# Standardizing workflows in imaging transcriptomics with the abagen toolbox

**DOI:** 10.1101/2021.07.08.451635

**Authors:** Ross D. Markello, Aurina Arnatkevičiūtė, Jean-Baptiste Poline, Ben D. Fulcher, Alex Fornito, Bratislav Misic

**Affiliations:** McConnell Brain Imaging Centre, Montréal Neurological Institute, McGill University, Montréal, Canada; Turner Institute for Brain and Mental Health, School of Psychological Sciences and Monash Biomedical Imaging, Monash University, Clayton, Australia; School of Physics, The University of Sydney, NSW 2006, Australia

**Keywords:** transcriptomics, neuroimaging, MRI, processing variability, software

## Abstract

Gene expression fundamentally shapes the structural and functional architecture of the human brain. Open-access transcriptomic datasets like the Allen Human Brain Atlas provide an unprecedented ability to examine these mechanisms *in vivo*; however, a lack of standardization across research groups has given rise to myriad processing pipelines for using these data. Here, we develop the `abagen` toolbox, an open-access software package for working with transcriptomic data, and use it to examine how methodological variability influences the outcomes of research using the Allen Human Brain Atlas. Applying three prototypical analyses to the outputs of 750,000 unique processing pipelines, we find that choice of pipeline has a large impact on research findings, with parameters commonly varied in the literature influencing correlations between derived gene expression and other imaging phenotypes by as much as *ρ* ≥ 1.0. Our results further reveal an ordering of parameter importance, with processing steps that influence gene normalization yielding the greatest impact on downstream statistical inferences and conclusions. The presented work and the development of the `abagen` toolbox lay the foundation for more standardized and systematic research in imaging transcriptomics, and will help to advance future understanding of the influence of gene expression in the human brain.

## INTRODUCTION

Technologies like magnetic resonance imaging (MRI) provide unique insights into macroscopic brain structure and function *in vivo*. Modern research increasingly emphasizes how microscale attributes, such as gene expression, influence these imaging-derived phenotypes (Arnatkeviciute et al. 2021, Arnatkevičiūtė et al. 2019, Fornito et al. 2019). Gene expression is particularly useful as it is a fundamental molecular phenotype that can be plausibly linked to the function of biological pathways (Seidlitz et al. 2018, Whitaker et al. 2016), protein synthesis (Zheng et al. 2019), receptor distributions (Beliveau et al. 2017, Deco et al. 2020, Nørgaard et al. 2021, Preller et al. 2018, Shine et al. 2019), and cell types (Anderson et al. 2020b, 2018, Gao et al. 2020, Hansen et al. 2021, Seidlitz et al. 2020). However, researchers looking to bridge these macro- and microscopic phenotypes must overcome multiple challenges. Although there are numerous technical and analytic considerations, one foundational issue is that acquiring high-quality transcriptomic data from the human brain is both costly and highly invasive, requiring budgets far greater than most typical neuroimaging studies and restrictive access to tissue from post-mortem donors or cranial surgical patients. As such, researchers must often rely on freely-available repositories of gene expression data.

There exist multiple open-access repositories for gene expression in the human brain, including BrainSpan (Kang et al. 2011, Miller et al. 2014) and PsychENCODE (Gandal et al. 2018, Li et al. 2018, Wang et al. 2018; among others: Darmanis et al. 2015, Lake et al. 2016, Sousa et al. 2017); however, these datasets generally provide relatively sparse anatomical coverage, limiting the types of analyses that can be performed. Thus, researchers who aim to compare transcriptomic expression with whole-brain imaging-derived phenotypes have primarily relied on the Allen Human Brain Atlas (AHBA; Hawrylycz et al. 2015, 2012). Initially released in 2010, the AHBA remains the most spatially comprehensive dataset of its kind. Derived from bulk microarray analysis of tissue samples obtained from six donors, the AHBA provides expression data for more than 20,000 genes across 3,702 brain areas in MRI-derived stereotactic space. With its superior resolution, the AHBA has significantly contributed to the emergence of the field of imaging transcriptomics (Fornito et al. 2019), enabling dozens of studies over the past decade examining relationships between gene expression and an array of macroscale imaging attributes, including cortical thickness (Shin et al. 2018), myelination (Burt et al. 2018), developmental brain maturation (Kirsch and Chechik 2016, Whitaker et al. 2016), structural brain networks (Arnatkevičiūtė et al. 2020, Romero-Garcia et al. 2018, Seidlitz et al. 2018), functional brain networks (Krienen et al. 2016, Richiardi et al. 2015, Vértes et al. 2016), and human cognition (Fox et al. 2014, Hansen et al. 2021). The AHBA has also highlighted the importance of whole-brain gene expression in neurological and psychiatric diseases, where it has become increasingly clear that transcriptional pathways play a critical role in shaping the broader dynamics of disease progression and emergent symptomatology (Anderson et al. 2020a, Henderson et al. 2019, McColgan et al. 2018, Morgan et al. 2019, Rittman et al. 2016, Romme et al. 2017, Shafiei et al. 2021, Vogel et al. 2020, Zheng et al. 2019).

Since its release, several software toolboxes have been developed to help researchers use transcriptional data from the AHBA (French and Paus 2015, Gorgolewski et al. 2015, Rittman et al. 2017, Rizzo et al. 2016); however, these tools often focus primarily on facilitating integration of the AHBA with neuroimaging data, offering limited if any functionality for modifying how the data are processed prior to analysis. Instead, a recent comprehensive review revealed that many research groups have opted to develop their own processing pipelines for the AHBA (Arnatkevičiūtė et al. 2019). Unfortunately, as there are no field-accepted standards for processing imaging transcriptomic data, the generated pipelines vary substantially across groups.

The extent to which such processing variability affects analytic outcomes from the AHBA remains unknown. Indeed, over the past decade neuroimaging research has shown that methodological variability can have broad influences on analyses using structural MRI (Bhagwat et al. 2021, Kharabian Masouleh et al. 2020), diffusion MRI (Maier-Hein et al. 2017, Oldham et al. 2020, Schilling et al. 2019), task fMRI (Botvinik-Nezer et al. 2020, Carp 2012), and resting-state fMRI (Ciric et al. 2017, Parkes et al. 2018). Although researchers are beginning to grapple with the consequences of this variability, the lack of baseline gene expression datasets against which to compare new results impedes the development of standardized practices. In these situations, some researchers have proposed performing “multiverse” analyses (Dragicevic et al. 2019, Steegen et al. 2016), wherein all possible permutations of data processing are analyzed and the full range of analytic results reported. Although such analyses can be computationally intensive, they offer a path to understand how processing choices impact statistical inferences and conclusions, and provide a mechanism by which to help researchers converge on an optimal pipeline.

Here, we comprehensively investigate how different processing choices influence the results of analyses using the AHBA. First, we develop an open-source Python toolbox, `abagen`, that collates all possible processing parameters into a set of turn-key workflows, optimized for flexibility and ease-of-use. We then use the toolbox to process the AHBA through approximately 750,000 unique pipelines. Across three prototypical imaging transcriptomic analyses, we examine whether and how these different processing options modify derived statistical estimates and quantify the relative importance of each option. Next, we replicate a curated set of processing pipelines from the literature to assess how previously-reported findings compare to the full range of potential outcomes observed across all examined pipelines. Finally, we end with a set of recommendations, integrated directly into the developed `abagen` toolbox, to promote standardized use of the AHBA in future work.

## RESULTS

We introduce the `abagen` toolbox, an open-access software package designed to streamline processing and preparation of the AHBA for integration with neuroimaging data (Markello et al. 2021, available at https://github.com/rmarkello/abagen). Supporting several workflows, `abagen` offers functionality for an array of analyses and has already been used in several peer-reviewed publications and preprints (Benkarim et al. 2020, Brown et al. 2021, Ding et al. 2021, Hansen et al. 2021, Lariviere et al. 2020, Martins et al. 2021, Park et al. 2021, 2020, Shafiei et al. 2021, 2020, Valk et al. 2021, Zhao et al. 2020). The primary workflow, used to generate regional gene expression matrices, integrates 17 distinct processing steps that have previously been employed by research groups throughout the published literature (Table 1). The following results use `abagen` to investigate how variable application of these processing steps can impact analyses of AHBA data.

**Table 1.** abagen pipeline options |. Overview of 17 options to be considered when processing the AHBA data. The *Choices* column indicates the number of parameters explored in the current report (numerator) and the total number of parameters possible for the given option (denominator). A denominator of *n* indicates a hypothetically near-infinite parameter space. The *Description* column gives a brief overview of the processing choice; for more detail refer to the relevant section in *Methods: Gene expression pipelines*.

| Option | Choices | Description |
| --- | --- | --- |
| Volumetric or surface atlas | 2/2 | Whether to use a volumetric or surface representation of the atlas |
| Individualized or group atlas | 1/2 | Whether to use individualized donor-specific atlases or a group-level atlas |
| Use non-linear MNI coordinates | 2/2 | Whether to use updated MNI coordinates provided by `alleninf` package |
| Mirror samples across L/R hemisphere | 3/4 | Whether to mirror (i.e., duplicate) samples across hemisphere boundary |
| Update probe-to-gene annotations | 2/2 | Whether to update probe annotations |
| Intensity-based filtering threshold | 3/$n$ | Threshold for intensity-based filtering of probes |
| Inter-areal similarity threshold | 1/$n$ | Threshold for removing samples with low inter-areal correspondence |
| Probe selection method | 6/8 | Method by which to select which probe(s) should represent a given gene |
| Donor-specific probe selection | 3/3 | How specified probe selection should integrate data from different donors |
| Missing data method | 2/3 | How to handle when brain regions are not assigned expression data |
| Sample-to-region matching tolerance | 3/$n$ | Distance tolerance for matching tissue samples to atlas brain regions |
| Sample normalization method | 3/10 | Method for normalizing tissue samples (across genes) |
| Gene normalization method | 3/10 | Method for normalizing genes (across tissue samples) |
| Normalize only matched samples | 2/2 | Whether to perform gene normalization for all versus matched samples |
| Normalizing discrete structures | 2/2 | Whether to perform gene normalization within structural classes |
| Sample-to-region combination method | 2/2 | Whether to aggregate tissue samples in regions within or across donors |
| Sample-to-region combination metric | 2/2 | Metric for aggregating tissue samples into atlas brain regions |

### Processing choices influence transcriptomic analyses

To understand how choices made during the processing of AHBA data impact downstream analyses, we enumerated 17 decision points (i.e., processing steps or options) that have been modified and used in the literature (Table 1). From these 17 steps we implemented 746,496 distinct processing pipelines, where each pipeline parcellated microarray expression from the AHBA with the Desikan-Killiany atlas (Desikan et al. 2006) to generate a unique brain region-by-gene expression matrix.

Analyses of expression data from the AHBA can be grouped into one of three broad classes (Fornito et al. 2019): correlated gene expression analyses, gene coexpression analyses, and regional gene expression analyses. Correlated gene expression analyses examine the correlation between brain regions across genes, yielding a symmetric region × region matrix (similar to a functional connectivity matrix). Gene co-expression analyses, on the other hand, examine the correlation between genes across brain regions, yielding a symmetric gene × gene matrix. Finally, regional gene expression analyses examine the expression patterns of specific genes or gene sets in relation to other imaging-derived phenotypes.

To examine how differences in processing choices may impact both the expression matrices generated from the different pipelines and derived statistical estimates we ran one analysis from each of these classes on the matrices generated by each processing pipeline. Notably, these analyses are either direct reproductions or variations of analyses that have been previously published (Arnatkeviĉiūtė et al. 2019, Burt et al. 2018, Hawrylycz et al. 2012, Oldham et al. 2008). Although there is no ground truth for any of these analyses, findings from previous work offer some context for interpreting the observed results (i.e., data from other species and other modalities; Lau et al. 2021). Nonetheless, we primarily focus on highlighting the potential variability resulting from different processing pipelines.

#### Correlated gene expression (CGE)

First, we separately correlated the rows of each expression matrix to generate symmetric region × region “correlated gene expression” matrices, indicating the similarity of gene expression profiles between different brain regions (Figure 1a). Previous work in other species has reliably observed that transcriptional similarity in the brain decays with increasing separation distance (Fulcher et al. 2019, Lau et al. 2021). This distance-dependent relationship is an expected feature due to the functional specialization of brain regions, and is consistent with other imaging-derived phenotypes in humans (Betzel and Bassett 2018, Goulas et al. 2019, Horvát et al. 2016, Mišić et al. 2014, Roberts et al. 2016, Shafiei et al. 2020). We assessed this relationship by extracting the upper triangle of the correlated gene expression matrices and correlating them with the upper triangle of a regional distance matrix, derived by computing the average Euclidean distance between brain regions in the Desikan-Killiany atlas (Fig. 1a, left panel). Although previous work has highlighted that this relationship is exponential (Arnatkevičiūtė et al. 2019), we computed the Spearman correlation as both statistics should exhibit similar variability across pipelines and the latter is less computationally expensive.

**Figure 1.**
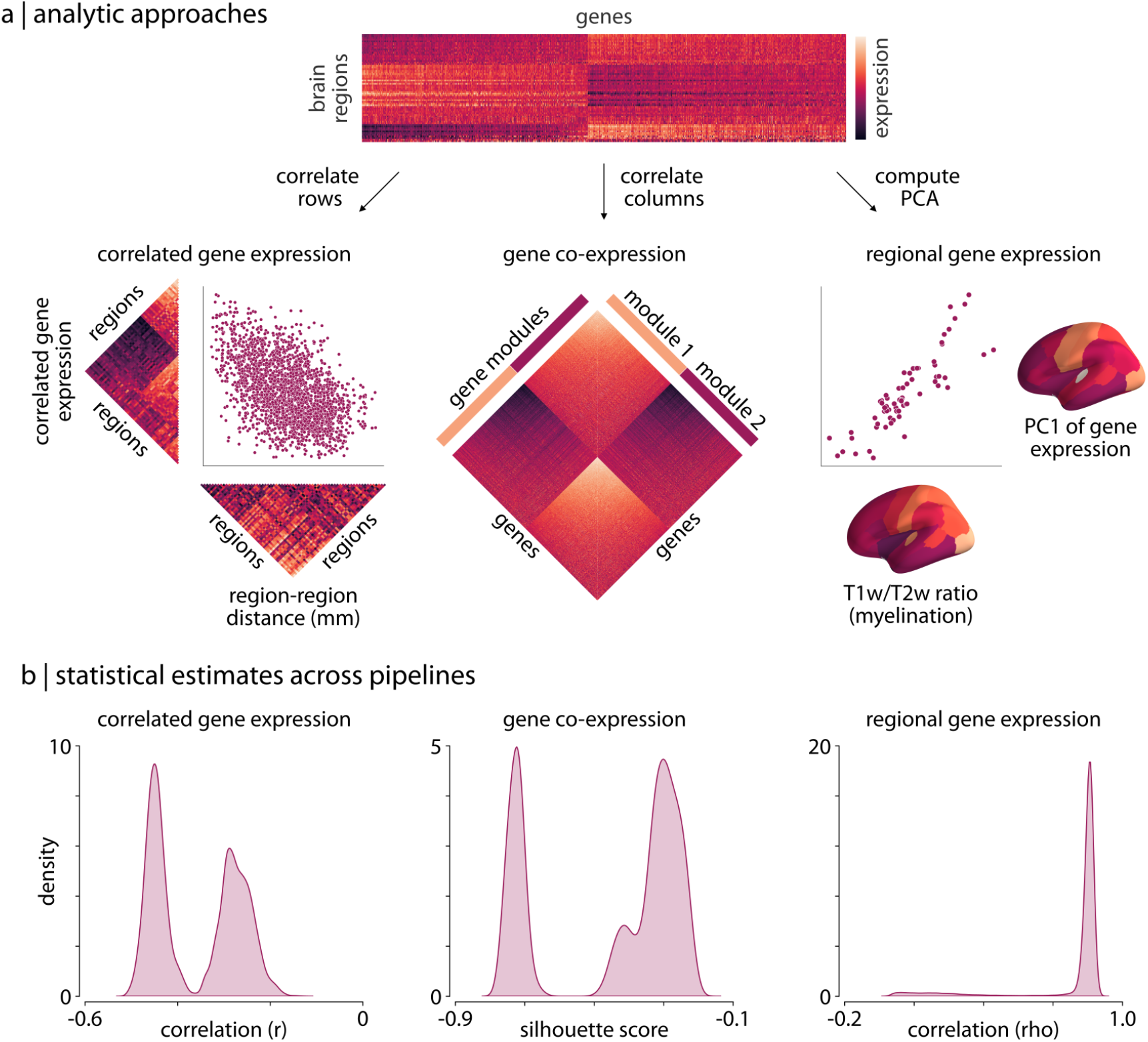
Processing choices influence transcriptomic analyses |. (a) Examples of the three analyses used to assess differences in gene expression matrices generated by transcriptomic pipelines. First row: a depiction of the region-by-gene expression matrix generated from one of the 746,946 tested processing pipelines. Second row, left: we compute the correlation between rows of each matrix to generate a symmetric region × region CGE matrix. We then compute the correlation between the upper triangle of this CGE matrix and the upper triangle of a regional distance matrix to examine the degree to which CGE decays with increasing distance between regions (Arnatkevičiūtė et al. 2019). Second row, middle: we compute the Euclidean distance between columns of each matrix to generate a gene × gene GCE matrix. We use previously defined functional gene communities (Oldham et al. 2008) to compute a silhouette score for this GCE matrix to investigate whether genes within a module have more similar patterns of spatial expression than genes between modules. Second row, right: the first principal component is extracted from the RGE matrix. We compute the correlation between this principal component and the whole-brain T1w/T2w ratio (Burt et al. 2018) to understand how closely these maps covary across the brain. (b) The full statistical distributions from each of the three analyses for all 746,496 pipelines. Left panel: Spearman correlation values, *ρ*, from the CGE analyses. Middle panel: silhouette scores from the GCE analyses. Right panel: Spearman correlation coefficients, *ρ*, from the RGE analyses. CGE: correlated gene expression; GCE: gene co-expression; RGE: regional gene expression.

#### Gene co-expression (GCE)

For the second type of analysis we separately correlated the columns of each expression matrix to generate gene × gene “co-expression” (GCE) matrices, indicating the similarity in spatial expression patterns between all pairs of genes (Figure 1a). A significant body of research has shown that genes tend to form functional communities, exhibiting synchronized expression patterns across space and time (Oldham et al. 2008), such that gene co-expression patterns tend to be more similar within than between such communities. Here, we obtained a set of gene community assignments derived for the brain from a previously studied human transcriptomic dataset (Oldham et al. 2008). We used these community assignments to calculate a silhouette score (Rousseeuw 1987) for the gene co-expression matrices generated by each pipeline, measuring how well these communities represented the derived co-expression patterns (Fig. 1a, middle panel).

#### Regional gene expression (RGE)

For the third type of transcriptomic analysis, we focused on regional correlations between gene expression measures and an MRI-derived phenotype. Our regional expression measure was defined by computing the first principal component of the region-by-gene expression matrix, representing the axis of maximum spatial variation of gene expression in the brain observed under a given AHBA processing pipeline. As gene expression fundamentally shapes the structure and function of the human brain, it is likely that this principal component may exhibit similar spatial variability to other imaging-derived measures. Recent work has highlighted that the T1w/T2w ratio is a robust phenotype that exhibits patterns of regional variation consistent with other microstructural and functional properties (Burt et al. 2018, Demirtaş et al. 2019, Fulcher et al. 2019, Gao et al. 2020). We therefore correlated the first principal component of gene expression with the wholebrain T1w/T2w ratio (Fig. 1a, right panel), measuring the extent to which these values covary across the cortex.

#### Pipeline distributions

Results from these three analyses reveal that choice of processing pipeline dramatically influences derived statistical estimates (i.e., the CGE-distance correlation, the gene co-expression silhouette score, and the spatial correlations between gene PC1 and whole-brain T1w/T2w ratio; Fig. 1b). We observe that all three of the generated distributions of statistical estimates across the 746,496 pipelines have wide ranges (correlated gene expression: [−0.51, −0.13]; gene co-expression: [−0.78, −0.18]; regional gene expression: [0.00, 0.90]) and are either bimodal (Fig. 1b, left/middle panels) or heavily skewed (Fig. 1b, right panel).

Since there is no ground truth for these analyses we cannot quantitatively assess whether some pipelines are more or less accurate than others. However, there is strong qualitative evidence to suggest that correlated gene expression should be lower between brain regions that are farther apart (Arnatkevičiūtė et al. 2019, Fulcher et al. 2019, Krienen et al. 2016, Lau et al. 2021, Richiardi et al. 2015). It is notable, then, that the distribution of distance-dependent estimates is so strongly bimodal (splitting at *r* ≈ −0.4), suggesting two very different perspectives on the size of this effect (Fig. 1a,b, left panels). As increasingly-detailed single-cell transcriptional data become available (e.g., Yao et al. 2021) we may be able to use these estimates to determine accuracy; for now, we simply note that even for this estimate with strong biological priors we see considerable variability.

Similar variability can be observed for the other two analyses. While all of the pipelines demonstrate relatively poor fit of gene communities to the derived gene co-expression matrices (refer to *Methods: Analytic approaches* for information on why this is not unexpected), we observe that a portion of the pipelines yield far worse correspondence (Fig. 1a,b, middle panels). Moreover, while the correlations between gene PC1 and whole-brain T1w/T2w ratio are largely consistent across pipelines, there are a small group of pipelines that yield correlations that deviate by *ρ* ≈ 1.0. Notably, the parameter choices for these pipelines are not pathological— that is, their use could be justified—and, as we discuss later (see *Results: Variability in parameter importance*), modifying just one parameter setting can yield changes in effect sizes within this range.

Collectively, we find that for all three of these analyses there is substantial variability in the statistical estimates generated by different processing pipelines, and this variability is large enough that, across pipelines, it has a meaningful difference in the potential inferences and conclusions that can be drawn.

### Variability in parameter importance

Next, we quantified the relative importance of different processing steps and parameters on our three derived statistical estimates. While researchers must ultimately make choices for each of the steps individually when processing AHBA data, we wanted to investigate whether unique choices have distinct influences. Moreover, which parameters are most important may differ based on the type of analysis performed.

We investigated parameter importance by calculating a distribution of difference scores for each parameter, measuring the extent to which changing each parameter— holding all other parameters constant—influences the derived statistical metrics from each of the three analyses. For example, given a processing parameter with two choices this procedure yielded a distribution of *N*/2 difference scores per analysis, where *N* is the total number of pipelines (i.e., 746,496/2 = 373,248). We averaged these distributions separately for each analysis to generate a single, summary “impact score” for each processing step, which we then rank-ordered independently for each analysis.

We find considerable agreement in which parameters are the most impactful across analyses (Fig. 2a): the most influential processing steps often involve procedures that influence the gene normalization process in some way (e.g., gene normalization method, normalizing only matched samples; Fig. 2b). On the other hand, among the least impactful parameters are choices concerning donor-specific probe selection and handling of missing data. It is worth noting that of the probe selection methods tested in the current manuscript (i.e., max intensity, correlation intensity, correlation variance, differential stability, RNAseq correlation, and averaging), three of the six all render the choice of donor-specific probe selection redundant. In other words, these three methods are mutually exclusive with choice of donor-specific probe selection, potentially confounding our ability to measure the real influence of this parameter. We also highlight that choice of atlas may influence the impact of missing data handling: since the Desikan-Killiany atlas is a relatively low-resolution atlas (68 nodes), expression matrices generated from the tested pipelines are missing, at most, data for two brain regions. It is possible that handling of missing data may be more important when higher-resolution parcellations are employed. That is, while some parameters do not appear to affect our results *in aggregate*, there are potentially specific research questions where these parameters could play an important and impactful role.

**Figure 2.**
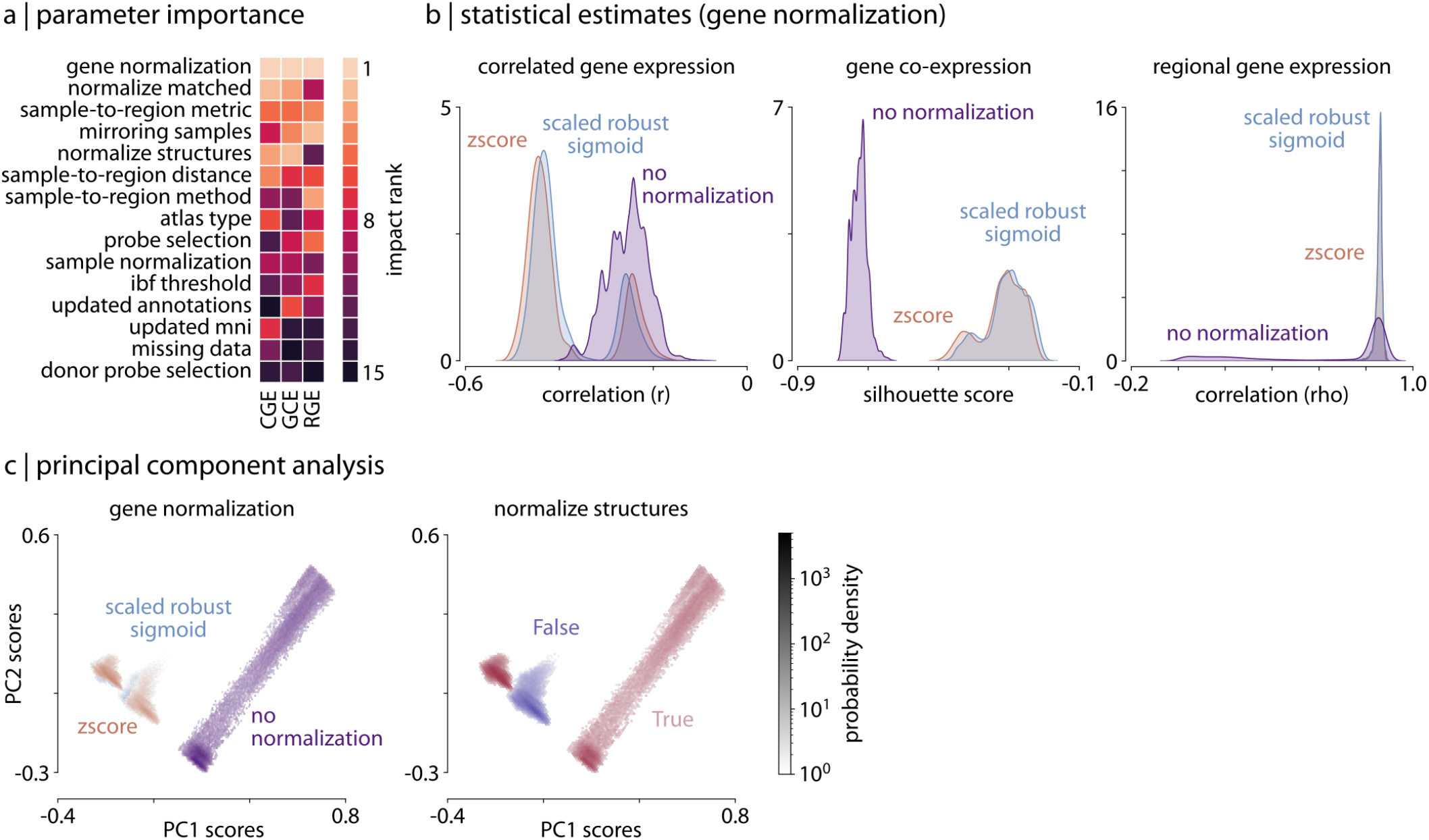
Parameter choice differentially impacts statistical estimates |. (a) Rank of the relative importance for each parameter (*y*-axis) across all three analyses (*x*-axis). Warmer colors indicate parameters that have a greater influence on statistical estimates. (b) Statistical distributions from the three analyses, shown as kernel density plots, separated by choice of gene normalization method (the most impactful parameter as shown in panel a). (c) Density plots of the statistical estimates for all 746,496 pipelines shown along the first two principal components, derived from the 746,496 (pipeline) x 3 (statistical estimates) matrix, representing how different the statistical estimates from each of the three analyses are relative to other pipelines. Left panel: pipelines are colored based on choice of gene normalization method, where each color represents 1/3 of the pipelines. Here, the pipelines in which no normalization was applied (purple) are distinguished from those in which some form of normalization was applied (blue and brown). Right panel: pipelines are colored based on whether gene normalization was performed within (True, red) or across (False, purple) structural classes (i.e., cortex, subcortex/brainstem, cerebellum; see *Methods: Gene expression pipelines* for more information).

To investigate those parameters that did play an influential role in the current analyses, we visualized their impact by examining the statistical distributions from each analysis separated by the different parameter choices (shown in Fig. 2b for gene normalization method). Dividing the distributions in this way highlights how strongly parameter choice can influence the outcomes of the analyses: for example, when no gene normalization is employed the resulting estimates are dramatically shifted from those generated by pipelines that employed some form of normalization (Fig. 2b; no normalization: blue distribution). Indeed, the bimodality and skew observed in the full statistical distributions for the analyses (Fig. 1b) is almost entirely explained by this single parameter choice.

To investigate more qualitative differences in how parameter choice influences the processing pipelines we performed a principal component analysis (PCA) on the matrix of statistical estimates from the three analyses (i.e., the 746,496 × 3 pipeline-by-analysis matrix). We extracted the first two principal components from the statistical estimate matrix (variance explained: PC1 = 70%, PC2 = 26%) and examined how pipeline scores were distributed along these axes (Fig. 2c). Delineating the distribution of pipelines based on parameter choice underscores how these options impact the separability of resulting statistical estimates. Reinforcing results presented above, we find that the choice of gene normalization method distinguishes the one-third of pipelines with no normalization (purple) from the remaining two-thirds that applied some form of normalization (blue and brown; Fig. 2c, left). It is clear from the distribution of pipelines, however, that other processing choices interact with this parameter. For example, plotting the pipelines by whether the gene normalization was performed separately on samples within each structural class (i.e., cerebral cortex, subcortex, cerebellum) rather than across all tissue samples further delineates the pipelines that applied gene normalization into two distinct clusters (Fig. 2c, right).

These results reveal how different processing steps are grouped in terms of their importance to analyses of the AHBA, with some groups demonstrating greater potential impact. Broadly, parameters modifying normalization are the most important, followed by parameters influencing how tissue samples are matched to brain regions, and finally parameters impacting probe selection. Moreover, we find that choices within each processing step do not all have an equivalent impact on derived estimates (i.e., performing no gene normalization has a much greater influence than choosing between the two other forms of normalization tested).

### Reproducing published analyses

The previous subsections demonstrate variability across the complete range of reasonable processing pipelines; however, many of these pipelines have not yet been used in practice. To investigate whether the subset of pipelines that have already been implemented in the published literature display similar variability, we used `abagen` to reproduce the processing procedures from nine peer-reviewed articles that (1) are highly-cited within the field, (2) highlight a wide range of processing options, and (3) sufficiently describe their processing pipelines such that they could be reproduced. We explored how different the gene expression values and statistical outcomes generated by these published pipelines were (Anderson et al. 2020b, 2018, Burt et al. 2018, French and Paus 2015, Hawrylycz et al. 2015, Krienen et al. 2016, Liu et al. 2020, Romero-Garcia et al. 2018, Whitaker et al. 2016). To ensure comparability, we standardized the choice of brain parcellation across pipelines, using the Desikan-Killiany atlas in all instances. The pipelines were used to generate nine region-by-gene expression matrices, which were then subjected to the same three analyses described previously.

In reproducing the pipelines we note important differences in processing parameter selection (Fig. 3a), and find that this variability results in slight discrepancies between gene expression values generated by the pipelines. For example, looking at the distribution of cortical somatostatin (SST), a gene discussed heavily in Anderson et al. (2020b) where it used as a proxy for somatostatin interneuron density (cf. Fulcher 2019), we observe some variation between pipelines (Fig. 3b,c). Although we find moderate consistency in the statistical estimates generated by the pipelines, there are important differences (ranges: correlated gene expression [−0.49, −0.28], gene co-expression [−0.70, −0.24], regional gene expression [0.34, 0.88]; Fig. 3c). One outlier is the single pipeline that did not appear to implement any form of gene normalization (French and Paus 2015), supporting earlier results demonstrating the importance of this processing step on downstream expression estimates. This is potentially notable as the processed expression data from this pipeline were made openly available and have been used in analyses by other researchers (e.g., Beliveau et al. 2017, Sepulcre et al. 2018).

**Figure 3.**
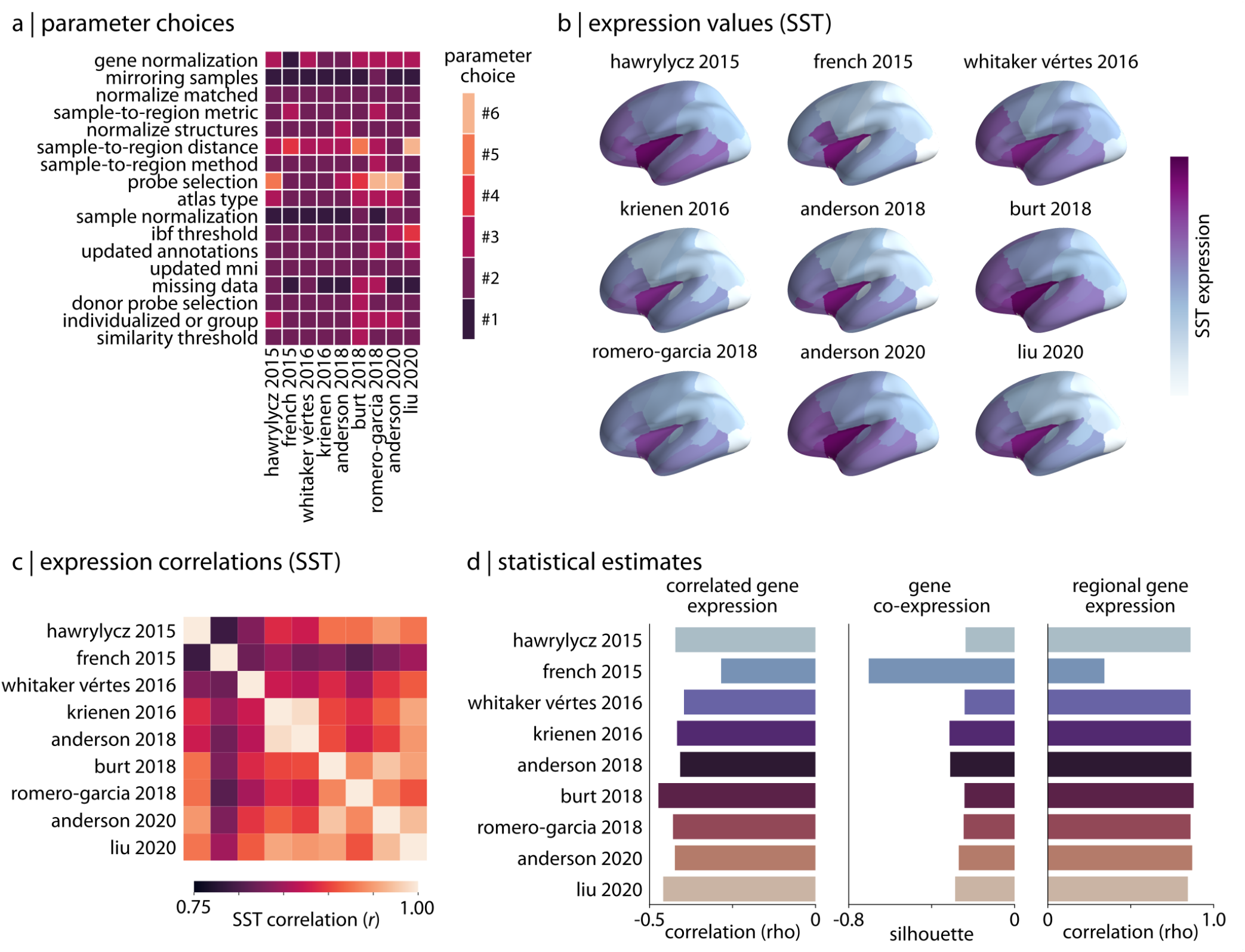
Reproducing published pipelines|. (a) Parameter choices used in the reproduction of published pipelines. Processing steps with categorical choices (e.g., gene normalization) were converted to numerical choices for display purposes only. These choices reflect the range of choices enumerated in Table 1. (b) Relative expression values of cortical somatostatin (SST) generated by each of the reproduced pipelines. Value ranges vary based on pipeline processing options. (c) The Pearson correlation between cortical somatostatin (SST) maps across the nine pipelines. (d) Statistical estimates from the three analyses described in *Methods: Analytic approaches* applied to expression data from each of the published pipelines.

Given that imaging transcriptomics is still relatively new and there has been limited work addressing best practices in the field (cf. Arnatkevičiūtė et al. 2019), these results stress the importance of standardization in use of the AHBA among research groups. Although variation in processing can ostensibly lead to similar inferences in specific analyses, even minor differences in processing choices consistently yield measurable discrepancies in derived expression data. Without proper standardization, these differences will compound and become more problematic as the field continues to grow.

### Standardized processing and reporting with the abagen toolbox

Across all of our analyses we find that choice of processing steps and parameters can have a strong influence on the statistical outcomes of research with the AHBA. Here, we briefly highlight features that we have integrated into the `abagen` toolbox to facilitate standardization in future research.

The `abagen` toolbox supports two use-case driven workflows: (1) a workflow that accepts an atlas and returns a parcellated, preprocessed regional gene expression matrix (Fig. 4a); and, (2) a workflow that accepts a mask and returns preprocessed expression data for all tissue samples within the mask (Fig. 4b). Workflows can be called via a single line of code from either the command line or Python terminal, and take approximately one minute to run with default settings using the Desikan-Killiany atlas. Although these workflows support the entire range of processing options that we assessed in the current manuscript (Fig. 4c), we have set the default options for all steps based on best practice recommendations developed in Arnatkevičiūtė et al. (2019) and further informed by the results presented above (see Table S1 for a full list).

**Figure 4.**
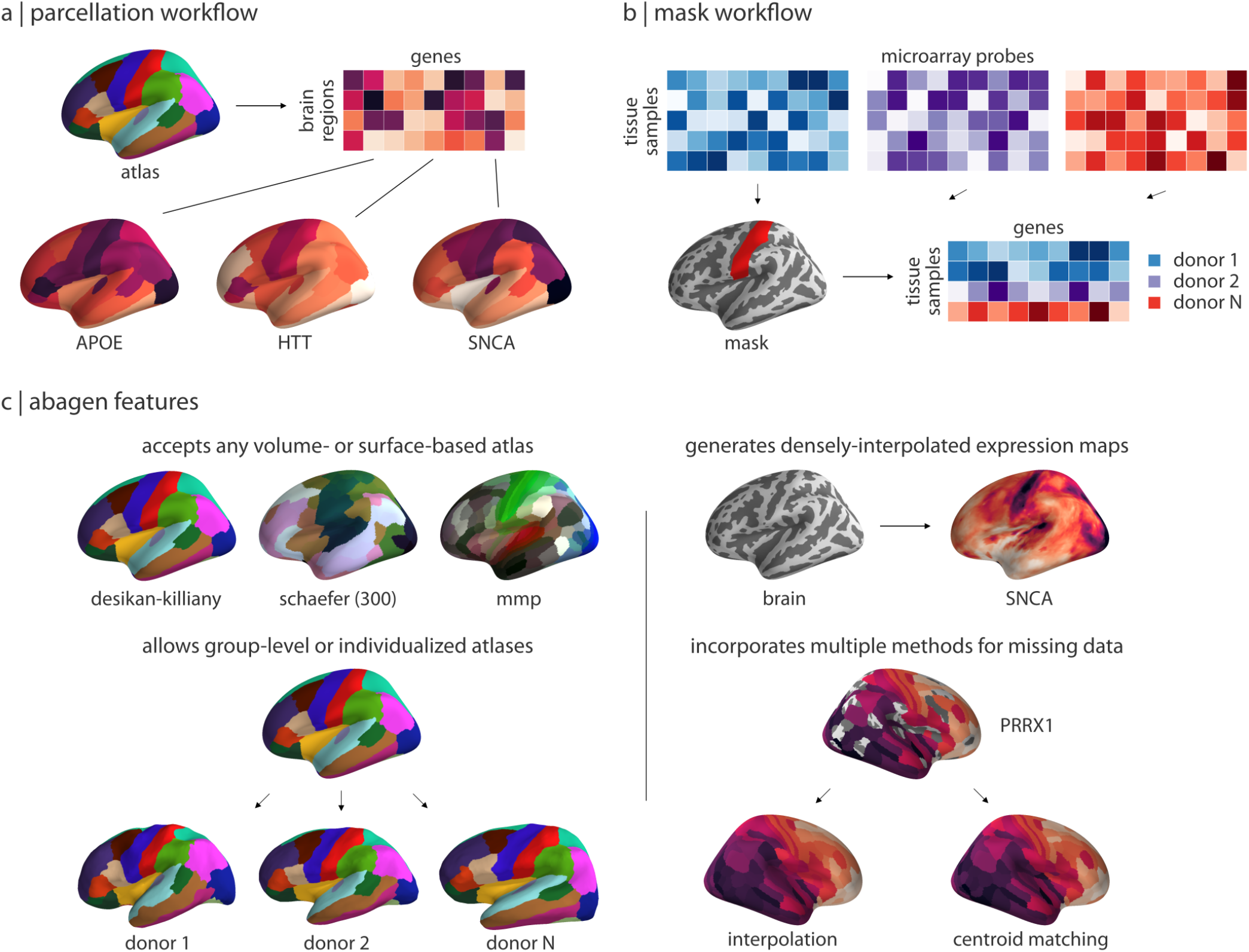
Workflows and features in the abagen toolbox |. (a) The primary workflow of `abagen`, used in the reported analyses, accepts a brain atlas and returns a parcellated brain-region-by-gene expression matrix. (b) An alternative `abagen` workflow accepts a regional mask and returns a processed tissue-sample-by-gene expression matrix, for all tissue samples from the six AHBA donors that fall within boundaries of the mask. (c) Examples of selected features from the `abagen` workflows and additional toolbox functionality. Top left: examples of some commonly-used atlases that can be employed with the parcellation workflow shown in panel (a). Bottom left: `abagen` can accept either standard atlases (i.e., in MNI space) or atlases defined in the space of the six individual donors from the AHBA. Top right: an additional workflow available in `abagen` can be used to generate densely-interpolated expression maps from AHBA data using a k-nearest neighbors interpolation algorithm. Bottom right: using high-resolution atlases in the parcellation workflow (panel a) may result in some parcels being assigned no expression data; `abagen` supports two methods for assigning values to such regions.

We believe the default settings in `abagen` will provide a reasonable starting point for researchers beginning to work with the AHBA; however, as we have continually noted, the appropriate choices for some parameters will vary based on research question. As such, to make it easier for researchers to report exactly what parameters they use, we have integrated an automated reporting mechanism into the `abagen` workflows (Fig. 5). The generated reports provide manuscript-ready step-by-step documentation describing all the processing done to the AHBA data in the workflow, and are licensed CC0 (https://creativecommons.org/share-your-work/public-domain/cc0/) so that they can be freely used without restriction.

**Figure 5.**
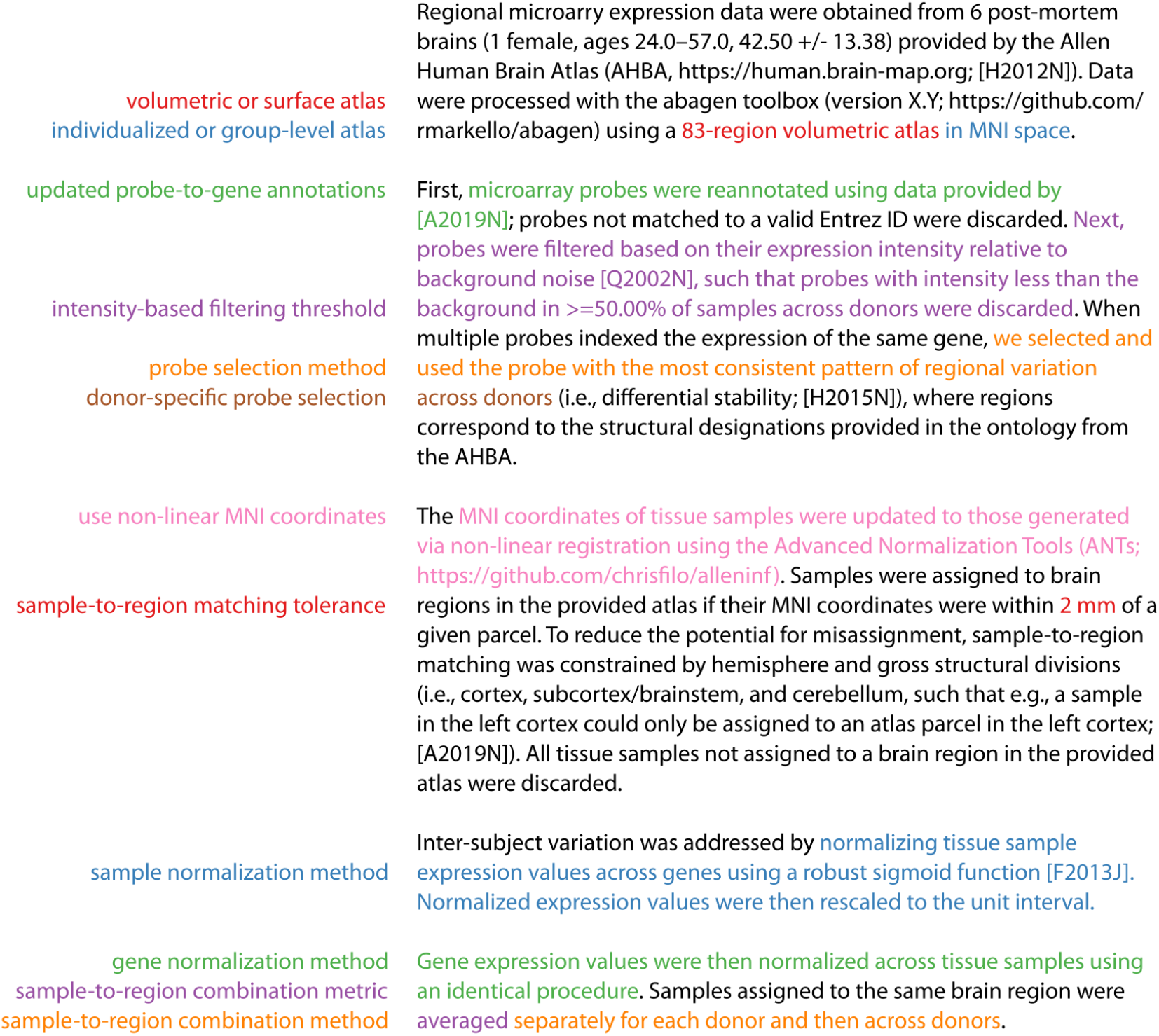
Annotated example abagen report |. Example of an automatically-generated methods section report from the `abagen` toolbox. Processing steps are shown on the left and the relevant methods text—which is updated when these steps are modified—is shown in the same font color on the right. Reports also include a formatted reference section and relevant equations; these are not shown here for conciseness. Note that some processing steps (e.g., normalizing within structures, missing data handling) are omitted here because they are not run by default (see Table S1).

Beyond its primary workflows, `abagen` has additional functionality for post-processing the AHBA data (e.g., removing distance-dependent effects from expression data, calculating differential stability estimates; Hawrylycz et al. 2015), and for accessing data from the companion Allen Mouse Brain Atlas (e.g., providing interfaces for querying the Allen Mouse API; https://mouse.brain-map.org/; Lein et al. 2007). Creation of the toolbox has followed best-practices in software development, including version control, continuous integration testing, and modular code design. `abagen` has already been successfully used in many peer-reviewed publications (Benkarim et al. 2020, Brown et al. 2021, Ding et al. 2021, Hansen et al. 2021, Lariviere et al. 2020, Martins et al. 2021, Park et al. 2021, 2020, Shafiei et al. 2021, 2020, Valk et al. 2021, Zhao et al. 2020), and we continue to integrate new features as community needs emerge. To encourage further use by new research groups we provide comprehensive documentation on installing and working with the `abagen` toolbox online (https://abagen.readthedocs.io/).

## DISCUSSION

In the present report we introduced the `abagen` toolbox, an open-source Python library for processing transcriptomic data. Using `abagen`, we conducted a comprehensive analysis examining whether and how different processing options modify statistical estimates derived from analyses using the AHBA. We investigated how processing pipelines used in the literature compare to those we tested, and provide recommendations for improving standardization and reporting of analyses using the AHBA, highlighting how the `abagen` toolbox can facilitate future developments in this space.

Testing nearly 750,000 unique processing pipelines, we find that choice of processing parameters can strongly influence statistical estimates derived from analyses of the AHBA, and that these choices interact with the type of analysis performed (Fig. 1). We observe significant variability with regard to which parameters are most influential, finding that procedures modifying gene expression normalization have a far greater impact on downstream analyses than other processing steps (Fig. 2). Looking to the literature, we reproduce nine pipelines from published articles and find that, despite notable inconsistencies in their processing choices, there is moderate consistency in their produced statistical estimates (Fig. 3). We demonstrate, however, that these summary estimates may obscure meaningful differences in gene expression values derived by the pipelines, cautioning researchers to be aware of how analytic choices may impact their findings.

Altogether, the present report provides a comprehensive assessment of how processing variability can impact analyses in the field of imaging transcriptomics. Our results demonstrate how researcher choices (or “researcher degrees of freedom”; Simmons et al. 2011) can play a meaningful role in analyses of the AHBA. However, these findings are not necessarily limited to the AHBA. Indeed, increasing reliance on open-access datasets has begun to reveal unique challenges associated with data reuse (Thompson et al. 2020). Improved standardization and reporting among research groups using (and re-using) openly-available datasets may help to mitigate some of these challenges. We believe that functionality in the `abagen` toolbox can support future researchers in overcoming these pitfalls and improve reproducibility in processing and analyzing AHBA data.

Our results also show that not all processing choices are equal: that is, we find a hierarchy of processing parameters, wherein procedures modifying gene normalization have the greatest impact on analyses, followed by steps more broadly influencing the matching of tissue samples to brain regions and finally by parameters that determine probe selection. Furthermore, we find that within processing steps certain parameter choices may lead to more reasonable statistical estimates. In particular, applying some form of gene normalization tends to improve the behavior of processed expression data when compared to instances in which no normalization is applied (Fig. 1), but there appear to be limited differences in the type of normalization used. Critically, these findings largely agree with previous recommendations developed by Arnatkevičiūtė et al. (2019), and we have chosen default parameter choices for `abagen` workflows accordingly.

More broadly, this work builds on increasing efforts to examine the importance of methodological choices and analytical flexibility in human neuroimaging research (Bhagwat et al. 2021, Botvinik-Nezer et al. 2020, Carp 2012, Ciric et al. 2017, Kharabian Masouleh et al. 2020, Maier-Hein et al. 2017, Oldham et al. 2020, Parkes et al. 2018, Schilling et al. 2019). Thankfully, emerging technical solutions have begun to tackle these issues via the development of tools that aim to abstract away sources of variation (e.g., fMRIPrep, Esteban et al. 2019; QSIPrep, Cieslak et al. 2020). While results from the present study reinforce the importance of methodological choices in research, `abagen` draws significant inspiration from these software packages in providing a set of tools designed to overcome such concerns when working with the AHBA.

While the AHBA dataset remains the only one of its kind, the `abagen` toolbox is designed to be used more broadly as similar datasets become available. That is, the preprocessing functions in `abagen` can be applied to other microarray expression datasets assuming, e.g., availability of stereotactic coordinates. As new imaging transcriptomic datasets are developed and become more widely-used, `abagen` functionality for creating standardized processing pipelines will only become more important. By developing the toolbox openly on GitHub (https://github.com/rmarkello/abagen), it is our hope that `abagen` can serve as a foundational, community tool for use in imaging transcriptomics research.

One consideration for future work on this topic is that the pipelines tested cover only a portion of the potential variability possible when processing AHBA data (Table 1). For example, a growing body of research has begun to examine how choice of brain parcellation may impact imaging analyses (e.g., Craddock et al. 2012, Markello and Misic 2021, Messé 2020, Thirion et al. 2014). While we only assessed processing pipelines using the Desikan-Killiany atlas, many other atlases have been used with the AHBA and it remains unclear how this variation may impact research findings. We also did not investigate whether donor-specific parcellations may impact analyses, a processing choice used in several published research findings (Anderson et al. 2020b, Burt et al. 2018, Romero-Garcia et al. 2018). Although there is significant evidence suggesting inter-individual variability in brain region definition (e.g., Dickie et al. 2018, Gordon et al. 2017, Kong et al. 2019), the process of generating individualized brain parcellations is fraught with methodological choices and requires careful data processing. Given the quality of the MRI data provided alongside the transcriptomic data in the AHBA—including important differences in scanning protocol and procedures between donors—creating donor-specific parcellations may be a large source of variability between pipelines.

Another limitation of the presented results is that we are unable to make categorical statements about which processing options are “best” for the AHBA. Unfortunately, there is no ground truth against which we can assess what the optimal set of processing parameters are, and encourage future work in this area to tackle this important problem. Moreover, the optimal set of processing parameters will very likely vary based on research question. Nonetheless, we offer two alternative solutions for researchers who want to continue using the AHBA data. First, similar to the current report, researchers can conduct a comprehensive analysis with the AHBA, running multiple processing pipelines and showing the entire distribution of generated statistical estimates; however, this process can be computationally prohibitive and may impair researchers’ abilities to interpret their findings (Steegen et al. 2016). A less costly alternative, then, is for the imaging transcriptomic research community to converge on a set of data-driven processing pipeline for the AHBA that can be used across research groups. We believe the `abagen` toolbox—with its comprehensive workflows, well-informed default parameter choices, and detailed documentation—can facilitate this process. While we acknowledge that some research groups may have strong reasons for wanting to use specific (i.e., non-default) processing choices, in these instances we urge clear and detailed reporting of the methods used—such as via the automated reporting functionality from the `abagen` toolbox.

Altogether, the current report highlights the problem of processing variability in analyses using the AHBA, impacting many research studies in the burgeoning field of imaging transcriptomics. We demonstrate how different processing options can influence statistical estimates of analyses relating data from the AHBA to imaging-derived phenotypes, and present the `abagen` toolbox as a promising potential solution to this issue.

## MATERIALS AND METHODS

### Code and data availability

All code used for data processing, analysis, and figure generation is available on GitHub (https://github.com/netneurolab/markello_transcriptome) and directly relies on the following open-source Python packages: IPython (Pérez and Granger 2007), Jupyter (Kluyver et al. 2016), Matplotlib (Hunter 2007), NiBabel (Brett et al. 2019), NumPy (Harris et al. 2020, Oliphant 2006, Van Der Walt et al. 2011), Pandas (McKinney et al. 2010), PySurfer (Waskom et al. 2020), Scikit-learn (Pedregosa et al. 2011), SciPy (Virtanen et al. 2020), and Seaborn (Waskom et al. 2018).

### Data

#### Allen Human Brain Atlas

The Allen Human Brain Atlas (AHBA) is an openaccess online resource containing whole-brain microarray gene expression data obtained from post-mortem tissue samples of six adult human donors (https://human.brain-map.org; Allen Institute for Brain Science 2013, Hawrylycz et al. 2012). Expression data for over 20,000 genes were sampled from 3,702 distinct tissue samples across the six donors (1 female, ages 24-57), providing the most spatially-comprehensive assay of gene expression in the human brain. Normalized microarray expression data were downloaded for all six donors; RNAseq data were downloaded for the two donors with relevant data.

#### Human Connectome Project

Group-averaged T1w/T2w (a proxy for intracortical myelin) data were downloaded from the S1200 release of the Human Connectome Project (HCP; Van Essen et al. 2013) and used without further processing.

#### Brain parcellations

All analyses were performed with the Desikan-Killiany atlas (DK; 68 cortical nodes), an anatomical parcellation generated by delineating regions based on gyral boundaries (Desikan et al. 2006). To explore the impact of volumetric-versus surface-based parcellations we used a version of the DK atlas in (1) volumetric MNI152, and (2) surface fsaverage5 space; both versions are provided directly with the `abagen` toolbox.

### The abagen toolbox

Source code for `abagen` is available on GitHub (https://github.com/rmarkello/abagen) and is provided under the three-clause BSD license (https://opensource.org/licenses/BSD-3-Clause). We have integrated `abagen` with Zenodo, which generates unique digital object identifiers (DOIs) for each new release of the toolbox (e.g., https://doi.org/10.5281/zenodo.3451463). Researchers can install `abagen` as a Python package via the PyPi repository (https://pypi.org/project/abagen/), and can access comprehensive online documentation via ReadTheDocs (https://abagen.readthedocs.io/).

### Gene expression pipelines

Most neuroimaging analyses using the AHBA must first convert the “raw” data into a pre-processed brain region-by-gene expression matrix. To investigate the extent to which different processing procedures might impact downstream analyses, we used `abagen` to modify 17 distinct processing steps in the generation of region-by-gene matrices from the original AHBA data, yielding 746,496 distinct pipelines. Here we describe in detail the 17 processing steps and respective methods for each option that we examined in our analyses (refer to Table 1 for a summary overview of these choices or refer to the `abagen` documentation for implementation details; https://abagen.readthedocs.io).

#### Volumetric or surface atlas

Aggregation of tissue samples from the AHBA into discrete brain regions requires researchers to supply an atlas (or parcellation). There are many brain atlases available for use, however they typically exist in one of two forms: defined (1) in 3D “volumetric” space, or (2) in “surface” space on a 2D representation of the cortical sheet. Many atlases can exist in both of these formats and so beyond the choice of parcellation, researchers must select which representation to use when processing AHBA samples. Choice of atlas may impact how many and which samples are matched to brain regions. In the current manuscript we examined a volume- and surface-based representation of the Desikan-Killiany atlas (see *Methods: Data*; Desikan et al. 2006). Note that both versions of the atlas used in the reported analyses are included with the “abagen” software distribution.

#### Individualized or group-level atlas

There is growing recognition that brain parcellations derived at the group level tend to obscure individual differences in anatomy or function (e.g., Dickie et al. 2018, Gordon et al. 2017, Kong et al. 2019). Researchers working with the AHBA have thus begun to generate donor-specific parcellations, using individualized atlases to match tissue samples to brain regions. The individualization process can vary dramatically depending on whether researchers are using volumetric or surface atlases and whether they are operating in “native” or standard (i.e., group) space. Because of the immense variability inherent to the individualization process itself, we opted not to explore this parameter in the current manuscript.

#### Use non-linear MNI coordinates

With its initial release the AHBA provided stereotactic coordinates for each tissue sample in MNI space (Collins et al. 1999, Fonov et al. 2011, 2009); however, two of the six donor brains were scanned *in cranio* and coordinates were derived using affine registrations to the MNI template, while the remaining four were scanned *ex vivo* and a non-linear registration was used to generate coordinates. More recently, Gorgolewski et al. (2014) used ANTS (Avants et al. 2011) to perform a standardized, manually-corrected non-linear diffeomorphic registration of all the donor brains to MNI space. Analyses collating tissue samples into distinct brain regions often rely on MNI coordinates to match samples to regions, and researchers must choose whether to use the original coordinates provided with the AHBA or the newer, non-linearly generated coordinates. In the current manuscript we assessed the impact of using (1) the original MNI coordinates and (2) the updated coordinates from Gorgolewski et al. (2014).

#### Mirror samples across left-right hemisphere

Only the first two donors included in the AHBA had tissue samples taken from the right hemisphere. Preliminary analyses of these data revealed minimal lateralization of microarray expression, and so samples were collected exclusively from the left hemisphere for the following four donors (Hawrylycz et al. 2015, 2012). This irregular sampling resulted in limited spatial coverage of expression in the right hemisphere; to resolve this, some researchers have opted to mirror existing tissue samples across the left-right hemisphere boundary (Romero-Garcia et al. 2018). Researchers must decide whether to perform sample mirroring, and, if so, whether they should mirror unilaterally (i.e., only right-to-left *or* left-to-right) or bilaterally (i.e., both right-to-left and left-to-right). In the current manuscript we assessed (1) no mirroring, (2) left-to-right mirroring, and (3) bilateral mirroring. The option for mirroring right-to-left was omitted as this is only useful when analyses selectively consider the left hemisphere, not the whole brain.

#### Update probe-to-gene annotations

The 60-base-pair probes used to assess microarray expression in the AHBA were annotated with their corresponding gene (or lack thereof) when the data were publicly released. However, as the human reference genome is updated these annotations become increasingly out-of-date. Thus, when researchers choose to use the AHBA data they must decide whether to use the original gene annotations or more recently-generated annotations. In the current manuscript we assessed using both the original annotations and those generated by Arnatkevičiūtė et al. (2019).

#### Intensity-based filtering threshold

Data from the AHBA are provided with information indicating whether the expression of each microarray probe exceeds the expression levels of background signal. Using this information, researchers can choose to perform an intensity-based filtering procedure wherein probes are only considered if their expression levels are greater than background across a specified percentage of tissue samples. In the current manuscript we considered three degrees of intensity-based filtering: (1) no filtering (all probes used), (2) 25% filtering (probes used if they exceeded background for more than 25% of all samples), and (3) median filtering (probes used if they exceeded background for more than 50% of all samples).

#### Inter-areal similarity threshold

The expression value of some tissue samples in the AHBA differ markedly from all other samples in the dataset. While this could be driven by real spatial variability in expression values throughout the brain, it is also possible that this variability is artifactual. Researchers can opt to asses the inter-areal similarity of tissue samples, quantifying those that differ from the rest by a given threshold, and remove them from consideration. To our knowledge this processing step has only been implemented in a single research study (Burt et al. 2018), and as such we do not consider it in the current manuscript.

#### Probe selection method

The probes used to measure microarray expression levels in the AHBA are often redundant; that is, there are frequently several probes indexing the same gene. Thus, at some point researchers must transition from measuring *probe* expression levels to measuring *gene* expression levels. Effectively, this means selecting from or condensing the redundant probes for each gene. There have been at least eight methods proposed in the literature for this process, including selecting a single probe with the (1) max intensity across samples, (2) max variance across samples, (3) highest loading on the first principal components across samples, (4) highest correlation to other probes (or max intensity across samples when only two probes exist), (5) highest correlation to other probes (or max variance across samples when only two probes exist), (6) highest differential stability across donors, (7) highest fidelity to simultaneously-acquired RNAseq data, or (8) simply averaging all probes indexing the same gene. In the current manuscript we only consider six of the most commonly-applied methods (i.e., 1, 4, 5, 6, 7, and 8); the other methods (i.e., 2 and 3) have only been reported in a single research study (Negi and Guda 2017 and Parkes et al. 2017, respectively) and as such we do not consider them.

#### Donor-specific probe selection

Probe selection (described above) often requires applying some selection criterion to gene expression levels across tissue samples. For these methods, the specified criterion can be measured across donors (i.e., aggregating tissues samples from donors) or independently for each donor. The latter case—performing probe selection independently for each donor—allows for two additional options: (1) using whichever probe is chosen for each donor, even if it differs from the other donors, or (2) using the most-commonly selected probe for all donors. In the current manuscript we considered all three of these options: (1) aggregating samples across donors, (2) performing probe selection independently for each donor, and (3) using the most commonly-selected probe across donors.

#### Missing data method

Due to the irregular spatial sampling of data in the AHBA some brain regions may not be assigned any corresponding microarray expression data. Researchers can opt to simply omit these regions from subsequent analyses; however, in some cases this is not desirable as the spatial distribution of the missing samples may not be random and discarding them may bias resulting estimates. Two options for handling missing data have been proposed in the literature, including filling missing regions with expression data from nearby regions (i.e., nearest-neighbors interpolation; Whitaker et al. 2016), or interpolating data in missing regions based on nearby samples (i.e., linear interpolation; Burt et al. 2018). In the current manuscript we tested two options: (1) omit brain regions with missing data entirely from subsequent analyses, and (2) fill missing data with expression values using nearest-neighbors interpolation. Linear interpolation has been sparingly used in the published literature (e.g., Burt et al. 2018, Romero-Garcia et al. 2018) and carries an increase in computational cost (approximately an order of magnitude higher than nearest neighbors interpolation); as such, we do not consider it in the current manuscript.

#### Sample-to-region matching tolerance

##### Volumetric atlases

While most tissue samples from the AHBA will fall directly within the brain regions delineated by most parcellations, some samples may fall outside the boundaries of these regions. Researchers can nonetheless choose to permit assigning these nearby samples to a given region, but will often set a distance threshold beyond which samples cannot be assigned. In the current manuscript we considered three distance tolerances: 0mm (i.e., samples must fall exactly within a region), 1mm, and 2mm.

##### Surface atlases

Because tissue samples from the AHBA are defined in volumetric space, matching them to parcels defined on a surface-based atlas requires different considerations than with volumetric atlases Notably, all samples will have non-zero distances from surface vertices; therefore, when matching to surface atlases distance thresholds are generally considered in terms of standard deviations (Burt et al. 2018; c.f., Anderson et al. 2020b). In this way all samples are matched to the surface and then those that are more than the specified standard deviation(s) above the mean away from the surface are excluded. In the current manuscript we tested three standard deviation distance tolerances: 0 s.d. (i.e., all samples farther than the average distance are excluded), 1 s.d., and 2 s.d..

#### Sample normalization method

Prior to aggregating microarray expression data across donors, researchers can optionally normalize the microarray expression data for each tissue sample across all represented genes (i.e., perform row-wise normalization). This procedure can account for between-sample differences in gene expression potentially driven by measurement errors. There is a number of techniques that have been proposed to normalize expression values; however, in the current manuscript we considered three normalization methods: (1) no normalization, (2) a z-score transform, and (3) a scaled robust sigmoid transform (Fulcher et al. 2013).

#### Gene normalization method

Prior to aggregating microarray expression data across donors, researchers can optionally normalize the microarray expression data for each represented gene across tissue samples (i.e., perform column-wise normalization). This procedure can account for interindividual (donor-specific) differences in gene expression data, which remain present in the AHBA despite batch corrections performed by the Allen Institute prior to releasing the data. In the current manuscript we considered three normalization methods: (1) no normalization, (2) a z-score transform, and (3) a scaled robust sigmoid transform (Fulcher et al. 2013).

#### Normalizing only matched samples

Due to choices in other processing steps (e.g., *Volume-or surface-based atlas, Sample-to-region matching tolerance* some tissue samples from the AHBA may not be assigned to any region in a given brain atlas. During gene normalization, where expression from each gene is normalized across tissue samples, researchers must decide whether to use (1) only those tissue samples matched to brain regions, or (2) the entire corpus of tissue samples, irrespective of whether they will be included in the final, processed regional expression matrix. In the current manuscript we consider both of these options.

#### Normalizing discrete structures

There is known variation in gene expression values between tissue samples taken from distinct structural classes (i.e., samples taken from neocortex may have different expression values than those from the brainstem). When performing gene normalization researchers can opt to normalize (1) across all samples irrespective of the structure from which they derive, or (2) independently for samples taken from different brain structures. Although the brain atlas used in the current manuscript represents only cortical parcels, this processing choice can interact with *Normalizing only matched samples* to impact resulting expression values and we therefore test both options.

Note that in the `abagen` toolbox structural classes are operationalized as: (1) cortex, (2) subcortex and brainstem, (3) cerebellum, and (4) white matter. Subcortex and brainstem are considered as one class because neuroanatomical delineation between these regions are widely contested and expression values in these regions tend to be more similar to one another than to other regions (i.e., data-driven clustering of samples tends to assign subcortical and brainstem samples together).

#### Sample-to-region combination method

Once tissue samples have been assigned to brain regions they need to be combined to generate a single expression profile; however, due to sampling differences between donors, some donors may have more tissue samples assigned to a given brain region than others. Thus, researchers must decide whether to aggregate samples (1) within each brain region independently for each donor and then across donors, or (2) simultaneously across all donors. In the latter case, donors with a higher number of samples matched to a region will contribute more to the expression profile of a given region (Arnatkevičiūtė et al. 2019). In the current manuscript we test both of these options.

#### Sample-to-region combination metric

When aggregating tissue samples into brain regions researchers must decide what aggregation metric they want to use. Although any statistical estimate could be considered, in practice an estimate of central tendency such as the mean expression values across tissue samples is most applicable. In the current manuscript we test aggregation with both the (1) mean and (2) median.

### Analytic approaches

Prototypical analyses relying on parcellated microarray expression data from the AHBA fall into three broad categories (Fornito et al. 2019):

1. *Correlated gene expression*: Examining the correlation between distinct brain regions across genes (i.e., using the region-by-region correlation matrix);
2. *Gene co-expression*: Examining the correlation between gene expression profiles across brain regions (i.e., using the gene-by-gene correlation matrix); or,
3. *Regional gene expression*: Examining the expression profile of one (or more) genes across brain regions (i.e., using selected columns of the region-by-gene expression matrix).

In order to examine the interaction between processing options and analytic method we performed one analysis from each of these three categories, described below, for every output of the 746,496 processing pipelines.

#### Correlated gene expression

Researchers have reliably found a relationship between correlated gene expression in the brain and the distance between brain regions: that is, brain regions that are farther away from one another tend to have less similar gene expression profiles (Arnatkevičiūtė et al. 2019, Krienen et al. 2016, Richiardi et al. 2017, 2015, Vértes et al. 2016). In order to examine the impact of processing choices on this relationship we computed the Spearman correlation between the upper triangle of the regional distance matrix (Euclidean distance between brain regions) and the upper triangle of each correlated gene expression matrix (Fig. 1a, left). Brain regions for which no gene expression data were available (dependent on pipeline options) were not included in the correlation. Note that this relationship is likely exponential (Arnatkevičiūtė et al. 2019); however, we calculated the Spearman coefficient as it is more computationally tractable and it should exhibit similar variability across pipelines.

#### Gene co-expression

Researchers have previously shown that gene expression in the brain tends to organize into functionally-defined communities or modules (Hawrylycz et al. 2012, Oldham et al. 2008). We examined the extent to which functional gene modules derived from a separate transcriptomic dataset (Oldham et al. 2008) mapped onto the gene co-expression matrices generated from the different processing pipelines. For each gene-by-gene matrix we calculated the silhouette score (Rousseeuw 1987) of the gene modules on a modified version of gene coexpression matrix (calculating Euclidean distance between genes instead of gene correlations; Fig. 1a, middle) via:

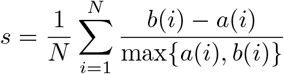

where *a(i)* is the average distance of a data point *i* to all other data points in the same cluster, *b(i)* is the mean distance of data point *i* to the nearest neighboring cluster, and N is the total number of data points. The final silhouette score s ranges from −1 to +1, where positive values indicate assortative and negative values indicate disassortative clusters.

Note that the original gene modules were defined using a weighted gene co-expression network analysis (WGCNA), which generally requires performing additional processing steps on the gene co-expression matrix. Since we used the raw gene co-expression matrix in the current analysis we expect lower silhouette scores than those reported in the initial manuscript where the gene communities were initially defined; however, the *variance* in scores between pipelines should not be significantly impacted by this choice.

#### Regional gene expression

Researchers recently highlighted how the principal component of gene expression in the brain closely mirrors the spatial variation observed in MRI-derived T1w/T2w measurements (typically used as a proxy for myelination; Burt et al. 2018). We examined whether this relationship was present across the outputs of the different pipelines, measuring the Spearman correlation between the T1w/T2w ratio and the first principal component of the regional gene expression matrix (Fig. 1a, right). Regional gene expression matrices were mean-centered prior to extraction of the principal component.

### Assessing pipeline impact

In order to examine the impact of each processing option on the resulting analyses we calculated a difference score, measuring the extent to which changing each option—holding all other options constant—influenced the derived metrics (i.e., correlation, silhouette score). When there were only two choices for a given option the impact was calculated as the absolute value of the difference between the two choices. When there were more than two choices and choices were ordinal (e.g., sample-to-region matching tolerance) the impact was calculated as the average of the absolute value of the difference between adjacent choices. When there were more than two choices and the choices were categorical (e.g., probe selection method) the impact was calculated as the average of the absolute value of the difference between all combinations of choices. These calculations yielded a distribution of “impact” estimates (i.e., change scores) for each processing option; we represented the final impact score for each processing option as the average of these distributions, taken independently for each of the three analyses. Impact estimates were rank-ordered (where the most impactful parameter was given a rank of one, the second most impactful a rank of two, and so on) to enable direct comparison across the different statistical estimates derived from the three analyses.

Researchers have previously shown that gene expression in the brain tends to organize into functionally-defined communities or modules (Hawrylycz et al. 2012, Oldham et al. 2008). We examined the extent to which functional gene modules derived from a separate transcriptomic dataset (Oldham et al. 2008) mapped onto the gene co-expression matrices generated from the different processing pipelines. For each gene-by-gene matrix we calculated the silhouette score (Rousseeuw 1987) of the gene modules on a modified version of gene coexpression matrix (calculating Euclidean distance between genes instead of gene correlations; Fig. 1a, middle) via:

### Pipeline dimensionality reduction

To investigate qualitative differences between the processing pipelines we performed a principal components analysis (PCA) on the matrix of estimates from the three statistical analyses (i.e., the 746,496 x 3 matrix). We mean-centered the columns of the matrix and extracted the first two principal components, examining how pipeline scores were distributed along these two components in relation to different processing options. These principal component highlight the closeness of the estimate generated by each pipeline along the dimensions of maximum statistical variation; that is, two pipelines that are closer together in the reduced-dimension space yielded more similar statistical estimates than two pipelines that are farther apart.

### Reproducing pipelines from the literature

Although all of the processing options explored in the current manuscript are reasonable or viable choices that researchers could make when preparing the AHBA for analysis, in reality these have not all been used in the published literature. In order to examine how pipelines used in the literature compared to those that we assessed, we selected nine articles that relied on data from the AHBA to support a primary research finding and reproduced their processing pipelines in `abagen` (Anderson et al. 2020b, 2018, Burt et al. 2018, French and Paus 2015, Hawrylycz et al. 2015, Krienen et al. 2016, Liu et al. 2020, Romero-Garcia et al. 2018, Whitaker et al. 2016). Note that these articles used a variety of parcellations and so to ensure comparability across pipelines we standardized this parameter, using the Desikan-Killiany atlas in all instances. One parameter that we did not assess in the pipelines explored in the current manuscript—whether to use individualized, donor-specific parcellations or a group-level atlas—was frequently varied in the published pipelines. Thus, when reproducing pipelines that called for individualized volumetric atlases we relied on the donor-specific Desikan-Killiany parcellations provided by Arnatkevičiūtė et al. (2019); when reproducing pipelines with individualized surface atlases we relied on the donor-specific Desikan-Killiany parcellations provided by Romero-Garcia et al. (2018).

As not all of the original manuscripts detailed the processing choices for each of the 17 steps in the `abagen` workflow, when specific parameter choices were omitted we either: (1) used the default setting if the parameter was required (e.g., using the mean for the “sample-to-region combination metric,” since all pipelines must combine samples to regions), or (2) omitted the processing step entirely if it is an optional step (e.g., not performing any gene normalization).

## ACKNOWLEDGEMENTS

We thank Vincent Bazinet, Elizabeth DuPre, Justine Hansen, Golia Shafiei, Laura Suárez, and Bertha Vázquez-Rodríguez for their comments and suggestions. This research was undertaken thanks in part to funding from the Canada First Research Excellence Fund, awarded to McGill University for the Healthy Brains for Healthy Lives initiative. This work was supported in part by funding provided by Brain Canada, in partnership with Health Canada, for the Canadian Open Neuroscience Platform initiative. RDM acknowledges support from the Fonds du Recherche Québec - Nature et Technologies and the Canadian Open Neuroscience Platform. BM acknowledges support from the Natural Sciences and Engineering Research Council of Canada (NSERC Discovery Grant RGPIN #017-04265) and from the Canada Research Chairs Program. AF was supported by the Sylvia and Charles Viertel Foundation and National Health and Medical Research Council (ID: 3274306). J-BP was partially funded by National Institutes of Health (NIH) NIH-NIBIB P41 EB019936 (ReproNim) NIH-NIMH R01 MH083320 (CANDIShare) and NIH RF1 MH120021 (NIDM), the National Institute Of Mental Health of the NIH under Award Number R01MH096906 (Neurosynth), and by Natural Sciences and Engineering Research Council of Canada (NSERC).

---

**Table S1.** Default abagen pipeline options |. The default settings for the 17 processing steps considered when processing the AHBA data with `abagen`. An entry of”—” indicates that this is a required, user-supplied parameter. A blank entry indicates that the processing step is not implemented by default. Refer to Table 1 and *Methods: Gene expression pipelines* for further details.

| *Option* | *Default* |
| --- | --- |
| Volumetric or surface atlas | — |
| Individualized or group atlas | — |
| Use non-linear MNI coordinates | True |
| Mirror samples across L/R hemisphere |  |
| Update probe-to-gene annotations | True |
| Intensity-based filtering threshold | 50% |
| Inter-areal similarity threshold |  |
| Probe selection method | differential stability |
| Donor-specific probe selection | aggregate |
| Missing data method |  |
| Sample-to-region matching tolerance | 2mm |
| Sample normalization method | scaled robust sigmoid |
| Gene normalization method | scaled robust sigmoid |
| Normalize only matched samples | True |
| Normalizing discrete structures | False |
| Sample-to-region combination method | donors |
| Sample-to-region combination metric | mean |

## References

Allen Institute for Brain Science (2013). Allen Human Brain Atlas online documentation. https://help.brain-map.org/display/humanbrain/Documentation.

Anderson, K. M., Collins, M. A., Chin, R., Ge, T., Rosenberg, M. D., and Holmes, A. J. (2020a). Transcriptional and imaging-genetic association of cortical interneurons, brain function, and schizophrenia risk. Nature Communications, 11(1):1–15.

Anderson, K. M., Collins, M. A., Kong, R., Fang, K., Li, J., He, T., Chekroud, A. M., Yeo, B. T., and Holmes, A. J. (2020b). Convergent molecular, cellular, and cortical neuroimaging signatures of major depressive disorder. Proceedings of the National Academy of Sciences, 117(40):25138–25149.

Anderson, K. M., Krienen, F. M., Choi, E. Y., Reinen, J. M., Yeo, B. T., and Holmes, A. J. (2018). Gene expression links functional networks across cortex and striatum. Nature Communications, 9(1):1–14.

Arnatkeviciute, A., Fulcher, B., Bellgrove, M., and Fornito, A. (2021). Where the genome meets the connectome: understanding how genes shape human brain connectivity. PsyArXiv.

Arnatkevičiūtė, A., Fulcher, B., Oldham, S., Tiego, J., Paquola, C., Gerring, Z., Aquino, K., Hawi, Z., Johnson, B., Ball, G., Klein, M., Deco, G., Franke, B., Bellgrove, M., and Fornito, A. (2020). Genetic influences on hub connectivity of the human connectome. bioRxiv.

Arnatkevičiūtė, A., Fulcher, B. D., and Fornito, A. (2019). A practical guide to linking brain-wide gene expression and neuroimaging data. NeuroImage, 189:353–367.

Avants, B. B., Tustison, N. J., Song, G., Cook, P. A., Klein, A., and Gee, J. C. (2011). A reproducible evaluation of ANTs similarity metric performance in brain image registration. NeuroImage, 54(3):2033–2044.

Beliveau, V., Ganz, M., Feng, L., Ozenne, B., Højgaard, L., Fisher, P. M., Svarer, C., Greve, D. N., and Knudsen, G. M. (2017). A high-resolution in vivo atlas of the human brain’s serotonin system. Journal of Neuroscience, 37(1):120–128.

Benkarim, O., Paquola, C., Park, B.-y., Hong, S.-J., Royer, J., de Wael, R. V., Larivière, S., Valk, S., Bzdok, D., Mottron, L., et al. (2020). Functional idiosyncrasy has a shared topography with group-level connectivity alterations in autism. bioRxiv.

Betzel, R. F. and Bassett, D. S. (2018). Specificity and robustness of long-distance connections in weighted, interareal connectomes. Proceedings of the National Academy of Sciences, 115(21):E4880–E4889.

Bhagwat, N., Barry, A., Dickie, E. W., Brown, S. T., Devenyi, G. A., Hatano, K., DuPre, E., Dagher, A., Chakravarty, M., Greenwood, C. M., et al. (2021). Understanding the impact of preprocessing pipelines on neuroimaging cortical surface analyses. GigaScience, 10(1):giaa155.

Botvinik-Nezer, R., Holzmeister, F., Camerer, C. F., Dreber, A., Huber, J., Johannesson, M., Kirchler, M., Iwanir, R., Mumford, J. A., Adcock, A., Avesani, P., Baczkowski, B., Bajracharya, A., Bakst, L., Ball, S., Barilari, M., Bault, N., Beaton, D., Beitner, J., Benoit, R., Berkers, R., Bhanji, J., Biswal, B., Bobadilla-Suarez, S., Bortolini, T., Bottenhorn, K., Bowring, A., Braem, S., Brooks, H., Brudner, E., Calderon, C., Camilleri, J., Castrellon, J., Cecchetti, L., Cieslik, E., Cole, Z., Collignon, O., Cox, R., Cunningham, W., Czoschke, S., Dadi, K., Davis, C., De Luca, A., Delgado, M., Demetriou, L., Dennison, J., Di, X., Dickie, E., Dobryakova, E., Donnat, C., Dukart, J., Duncan, N. W., Durnez, J., Eed, A., Eickhoff, S., Erhart, A., Fontanesi, L., Fricke, G. M., Galvan, A., Gau, R., Genon, S., Glatard, T., Glerean, E., Goeman, J., Golowin, S., González-García, C., Gorgolewski, K., Grady, C., Green, M., Guassi Moreira, J., Guest, O., Hakimi, S., Hamilton, J. P., Hancock, R., Handjaras, G., Harry, B., Hawco, C., Herholz, P., Herman, G., Heunis, S., Hoffstaedter, F., Hogeveen, J., Holmes, S., Hu, C.-P., Huettel, S., Hughes, M., Iacovella, V., Iordan, A., Isager, P., Isik, A. I., Jahn, A., Johnson, M., Johnstone, T., Joseph, M., Juliano, A., Kable, J., Kassinopoulos, M., Koba, C., Kong, X.-Z., Koscik, T., Kucukboyaci, N. E., Kuhl, B., Kupek, S., Laird, A., Lamm, C., Langner, R., Lauharatanahirun, N., Lee, H., Lee, S., Leemans, A., Leo, A., Lesage, E., Li, F., Li, M., Lim, P. C., Lintz, E., Liphardt, S., Losecaat Vermeer, A., Love, B., Mack, M., Malpica, N., Marins, T., Maumet, C., McDonald, K., McGuire, J., Melero, H., Méndez Leal, A., Meyer, B., Meyer, K., Mihai, P., Mitsis, G., Moll, J., Nielson, D., Nilsonne, G., Notter, M., Olivetti, E., Onicas, A., Papale, P., Patil, K., Peelle, J. E., Pérez, A., Pischedda, D., Poline, J.-B., Prystauka, Y., Ray, S., Reuter-Lorenz, P., Reynolds, R., Ricciardi, E., Rieck, J., Rodriguez-Thompson, A., Romyn, A., Salo, T., Samanez-Larkin, G., Sanz-Morales, E., Schlichting, M., Schultz, D., Shen, Q., Sheridan, M., Shiguang, F., Silvers, J., Skagerlund, K., Smith, A., Smith, D., Sokol-Hessner, P., Steinkamp, S., Tashjian, S., Thirion, B., Thorp, J., Tinghög, G., Tisdall, L., Tompson, S., Toro-Serey, C., Torre, J., Tozzi, L., Truong, V., Turella, L., van’t Veer, A. E., Verguts, T., Vettel, J., Vijayarajah, S., Vo, K., Wall, M., Weeda, W. D., Weis, S., White, D., Wisniewski, D., Xifra-Porxas, A., Yearling, E., Yoon, S., Yuan, R., Yuen, K., Zhang, L., Zhang, X., Zosky, J., Nichols, T. E., Poldrack, R. A., and Schonberg, T. (2020). Variability in the analysis of a single neuroimaging dataset by many teams. Nature, 582:84–88.

Brett, M., Markiewicz, C. J., Hanke, M., Côté, M.-A., Cipollini, B., McCarthy, P., Cheng, C. P., Halchenko, Y. O., Cottaar, M., Ghosh, S., and et al. (2019). nipy/nibabel. Zenodo, doi:10.5281/zenodo.591597.

Brown, J. A., Lee, A. J., Pasquini, L., and Seeley, W. W. (2021). A dynamic gradient architecture generates brain activity states. bioRxiv, pages 2020–08.

Burt, J. B., Demirtaş, M., Eckner, W. J., Navejar, N. M., Ji, J. L., Martin, W. J., Bernacchia, A., Anticevic, A., and Murray, J. D. (2018). Hierarchy of transcriptomic specialization across human cortex captured by structural neuroimaging topography. Nature neuroscience, 21(9):1251–1259.

Carp, J. (2012). On the plurality of (methodological) worlds: estimating the analytic flexibility of fMRI experiments. Frontiers in Neuroscience, 6:149.

Cieslak, M., Cook, P. A., He, X., Yeh, F.-C., Dhollander, T., Adebimpe, A., Aguirre, G. K., Bassett, D. S., Betzel, R. F., Bourque, J., et al. (2020). QSIPrep: An integrative platform for preprocessing and reconstructing diffusion MRI. bioRxiv.

Ciric, R., Wolf, D. H., Power, J. D., Roalf, D. R., Baum, G. L., Ruparel, K., Shinohara, R. T., Elliott, M. A., Eickhoff, S. B., Davatzikos, C., et al. (2017). Benchmarking of participant-level confound regression strategies for the control of motion artifact in studies of functional connectivity. NeuroImage, 154:174–187.

Collins, D. L., Zijdenbos, A. P., Baaré, W. F., and Evans, A. C. (1999). ANIMAL+INSECT: improved cortical structure segmentation. In Biennial International Conference on Information Processing in Medical Imaging, pages 210–223. Springer.

Craddock, R. C., James, G. A., Holtzheimer III, P. E., Hu, X. P., and Mayberg, H. S. (2012). A whole brain fMRI atlas generated via spatially constrained spectral clustering. Human Brain Mapping, 33(8):1914–1928.

Darmanis, S., Sloan, S. A., Zhang, Y., Enge, M., Caneda, C., Shuer, L. M., Gephart, M. G. H., Barres, B. A., and Quake, S. R. (2015). A survey of human brain transcriptome diversity at the single cell level. Proceedings of the National Academy of Sciences, 112(23):7285–7290.

Deco, G., Aquino, K. M., Arnatkeviciute, A., Oldham, S., Sabaroedin, K., Rogasch, N. C., Kringelbach, M. L., and Fornito, A. (2020). Dynamical consequences of regional heterogeneity in the brains transcriptional landscape. bioRxiv.

Demirtaş, M., Burt, J. B., Helmer, M., Ji, J. L., Adkinson, B. D., Glasser, M. F., Van Essen, D. C., Sotiropoulos, S. N., Anticevic, A., and Murray, J. D. (2019). Hierarchical heterogeneity across human cortex shapes large-scale neural dynamics. Neuron, 101(6):1181–1194.

Desikan, R. S., Ségonne, F., Fischl, B., Quinn, B. T., Dickerson, B. C., Blacker, D., Buckner, R. L., Dale, A. M., Maguire, R. P., Hyman, B. T., et al. (2006). An automated labeling system for subdividing the human cerebral cortex on MRI scans into gyral based regions of interest. NeuroImage, 31(3):968–980.

Dickie, E. W., Ameis, S. H., Shahab, S., Calarco, N., Smith, D. E., Miranda, D., Viviano, J. D., and Voineskos, A. N. (2018). Personalized intrinsic network topography mapping and functional connectivity deficits in autism spectrum disorder. Biological Psychiatry, 84(4):278–286.

Ding, Y., Zhao, K., Che, T., Du, K., Sun, H., Liu, S., Zheng, Y., Li, S., Liu, B., Liu, Y., et al. (2021). Quantitative radiomic features as new biomarkers for alzheimer’s disease: An amyloid pet study. Cerebral Cortex.

Dragicevic, P., Jansen, Y., Sarma, A., Kay, M., and Chevalier, F. (2019). Increasing the transparency of research papers with explorable multiverse analyses. In Proceedings of the 2019 CHI Conference on Human Factors in Computing Systems, pages 1–15.

Esteban, O., Markiewicz, C. J., Blair, R. W., Moodie, C. A., Isik, A. I., Erramuzpe, A., Kent, J. D., Goncalves, M., DuPre, E., Snyder, M., et al. (2019). fMRIPrep: a robust preprocessing pipeline for functional MRI. Nature Methods, 16(1):111–116.

Fonov, V., Evans, A. C., Botteron, K., Almli, C. R., McKinstry, R. C., Collins, D. L., Group, B. D. C., et al. (2011). Unbiased average age-appropriate atlases for pediatric studies. NeuroImage, 54(1):313–327.

Fonov, V. S., Evans, A. C., McKinstry, R. C., Almli, C., and Collins, D. (2009). Unbiased nonlinear average age-appropriate brain templates from birth to adulthood. NeuroImage, 47:S102.

Fornito, A., Arnatkevičiūtė, A., and Fulcher, B. D. (2019). Bridging the gap between connectome and transcriptome. Trends in Cognitive Sciences, 23(1):34–50.

Fox, A. S., Chang, L. J., Gorgolewski, K. J., and Yarkoni, T. (2014). Bridging psychology and genetics using large-scale spatial analysis of neuroimaging and neurogenetic data. bioRxiv, page 012310.

French, L. and Paus, T. (2015). A Freesurfer view of the cortical transcriptome generated from the Allen Human Brain Atlas. Frontiers in Neuroscience, 9:323.

Fulcher, B. D. (2019). Discovering conserved properties of brain organization through multimodal integration and interspecies comparison. Journal of Experimental Neuroscience, 13:1179069519862047.

Fulcher, B. D., Little, M. A., and Jones, N. S. (2013). Highly comparative time-series analysis: the empirical structure of time series and their methods. Journal of the Royal Society Interface, 10(83):20130048.

Fulcher, B. D., Murray, J. D., Zerbi, V., and Wang, X.-J. (2019). Multimodal gradients across mouse cortex. Proceedings of the National Academy of Sciences, 116(10):4689–4695.

Gandal, M. J., Zhang, P., Hadjimichael, E., Walker, R. L., Chen, C., Liu, S., Won, H., Van Bakel, H., Varghese, M., Wang, Y., et al. (2018). Transcriptome-wide isoform-level dysregulation in asd, schizophrenia, and bipolar disorder. Science, 362(6420).

Gao, R., van den Brink, R. L., Pfeffer, T., and Voytek, B. (2020). Neuronal timescales are functionally dynamic and shaped by cortical microarchitecture. Elife, 9:e61277.

Gordon, E. M., Laumann, T. O., Gilmore, A. W., Newbold, D. J., Greene, D. J., Berg, J. J., Ortega, M., Hoyt-Drazen, C., Gratton, C., Sun, H., et al. (2017). Precision functional mapping of individual human brains. Neuron, 95(4):791–807.

Gorgolewski, K. J., Fox, A. S., Chang, L., Schäfer, A., Arélin, K., Burmann, I., Sacher, J., and Margulies, D. S. (2014). Tight fitting genes: finding relations between statistical maps and gene expression patterns. F1000Research, 5.

Gorgolewski, K. J., Varoquaux, G., Rivera, G., Schwarz, Y., Ghosh, S. S., Maumet, C., Sochat, V. V., Nichols, T. E., Poldrack, R. A., Poline, J.-B., et al. (2015). NeuroVault.org: a web-based repository for collecting and sharing unthresholded statistical maps of the human brain. Frontiers in Neuroinformatics, 9:8.

Goulas, A., Betzel, R. F., and Hilgetag, C. C. (2019). Spatiotemporal ontogeny of brain wiring. Science Advances, 5(6):eaav9694.

Hansen, J. Y., Markello, R. D., Vogel, J. W., Seidlitz, J., Bzdok, D., and Misic, B. (2021). Mapping gene transcription and neurocognition across human neocortex. Nature Human Behaviour, pages 1–11.

Harris, C. R., Millman, K. J., van der Walt, S. J., Gommers, R., Virtanen, P., Cournapeau, D., Wieser, E., Taylor, J., Berg, S., Smith, N. J., et al. (2020). Array programming with NumPy. Nature, 585(7825):357–362.

Hawrylycz, M., Miller, J. A., Menon, V., Feng, D., Dolbeare, T., Guillozet-Bongaarts, A. L., Jegga, A. G., Aronow, B. J., Lee, C.-K., Bernard, A., et al. (2015). Canonical genetic signatures of the adult human brain. Nature Neuroscience, 18(12):1832.

Hawrylycz, M. J., Lein, E. S., Guillozet-Bongaarts, A. L., Shen, E. H., Ng, L., Miller, J. A., Van De Lagemaat, L. N., Smith, K. A., Ebbert, A., Riley, Z. L., et al. (2012). An anatomically comprehensive atlas of the adult human brain transcriptome. Nature, 489(7416):391.

Henderson, M. X., Cornblath, E. J., Darwich, A., Zhang, B., Brown, H., Gathagan, R. J., Sandler, R. M., Bassett, D. S., Trojanowski, J. Q., and Lee, V. M. (2019). Spread of *α*-synuclein pathology through the brain connectome is modulated by selective vulnerability and predicted by network analysis. Nature Neuroscience, 22(8):1248–1257.

Horvát, S., Gămănut, R., Ercsey-Ravasz, M., Magrou, L., Gămănut, B., Van Essen, D. C., Burkhalter, A., Knoblauch, K., Toroczkai, Z., and Kennedy, H. (2016). Spatial embedding and wiring cost constrain the functional layout of the cortical network of rodents and primates. PLoS Biology, 14(7):e1002512.

Hunter, J. D. (2007). Matplotlib: A 2D graphics environment. Computing in Science & Engineering, 9(3):90–95.

Kang, H. J., Kawasawa, Y. I., Cheng, F., Zhu, Y., Xu, X., Li, M., Sousa, A. M., Pletikos, M., Meyer, K. A., Sedmak, G., et al. (2011). Spatio-temporal transcriptome of the human brain. Nature, 478(7370):483–489.

Kharabian Masouleh, S., Eickhoff, S. B., Zeighami, Y., Lewis, L. B., Dahnke, R., Gaser, C., Chouinard-Decorte, F., Lepage, C., Scholtens, L. H., Hoffstaedter, F., et al. (2020). Influence of processing pipeline on cortical thickness measurement. Cerebral Cortex, 30(9):5014–5027.

Kirsch, L. and Chechik, G. (2016). On expression patterns and developmental origin of human brain regions. PLoS computational biology, 12(8):e1005064.

Kluyver, T., Ragan-Kelley, B., Pérez, F., Granger, B. E., Bussonnier, M., Frederic, J., Kelley, K., Hamrick, J. B., Grout, J., Corlay, S., et al. (2016). Jupyter Notebooks-A publishing format for reproducible computational workflows. In Loizides, F. and Scmidt, B., editors, Positioning and Power in Academic Publishing: Players, Agents and Agendas, pages 87–90. IOS Press.

Kong, R., Li, J., Orban, C., Sabuncu, M. R., Liu, H., Schaefer, A., Sun, N., Zuo, X.-N., Holmes, A. J., Eickhoff, S. B., et al. (2019). Spatial topography of individual-specific cortical networks predicts human cognition, personality, and emotion. Cerebral Cortex, 29(6):2533–2551.

Krienen, F. M., Yeo, B. T., Ge, T., Buckner, R. L., and Sherwood, C. C. (2016). Transcriptional profiles of supragranular-enriched genes associate with corticocortical network architecture in the human brain. Proceedings of the National Academy of Sciences, 113(4):E469–E478.

Lake, B. B., Ai, R., Kaeser, G. E., Salathia, N. S., Yung, Y. C., Liu, R., Wildberg, A., Gao, D., Fung, H.-L., Chen, S., et al. (2016). Neuronal subtypes and diversity revealed by single-nucleus rna sequencing of the human brain. Science, 352(6293):1586–1590.

Lariviere, S., Paquola, C., Park, B.-y., Royer, J., Wang, Y., Benkarim, O., de Wael, R. V., Valk, S. L., Thomopoulos, S. I., Kirschner, M., et al. (2020). The ENIGMA toolbox: Crossdisorder integration and multiscale neural contextualization of multisite neuroimaging datasets. bioRxiv.

Lau, H. Y. G., Fornito, A., and Fulcher, B. D. (2021). Scaling of gene transcriptional gradients with brain size across mouse development. NeuroImage, 224:117395.

Lein, E. S., Hawrylycz, M. J., Ao, N., Ayres, M., Bensinger, A., Bernard, A., Boe, A. F., Boguski, M. S., Brockway, K. S., Byrnes, E. J., et al. (2007). Genome-wide atlas of gene expression in the adult mouse brain. Nature, 445(7124):168–176.

Li, M., Santpere, G., Kawasawa, Y. I., Evgrafov, O. V., Gulden, F. O., Pochareddy, S., Sunkin, S. M., Li, Z., Shin, Y., Zhu, Y., et al. (2018). Integrative functional genomic analysis of human brain development and neuropsychiatric risks. Science, 362(6420).

Liu, J., Xia, M., Wang, X., Liao, X., and He, Y. (2020). The spatial organization of the chronnectome associates with cortical hierarchy and transcriptional profiles in the human brain. Neuroimage, 222:117296.

Maier-Hein, K. H., Neher, P. F., Houde, J.-C., Côté, M.-A., Garyfallidis, E., Zhong, J., Chamberland, M., Yeh, F.-C., Lin, Y.-C., Ji, Q., et al. (2017). The challenge of mapping the human connectome based on diffusion tractography. Nature communications, 8(1):1–13.

Markello, R., Shafiei, G., Zheng, Y.-Q., and Mišić, B. (2021). rmarkello/abagen. Zenodo, doi:10.5281/zenodo.3451463.

Markello, R. D. and Misic, B. (2021). Comparing spatial null models for brain maps. NeuroImage, page 118052.

Martins, D., Dipasquale, O., Veronese, M., Turkheimer, F. E., Loggia, M., McMahon, S., and Williams, S. C. (2021). Transcriptional and cellular signatures of cortical morphometric similarity remodelling in chronic pain. bioRxiv.

McColgan, P., Gregory, S., Seunarine, K. K., Razi, A., Papoutsi, M., Johnson, E., Durr, A., Roos, R. A., Leavitt, B. R., Holmans, P., et al. (2018). Brain regions showing white matter loss in huntington’s disease are enriched for synaptic and metabolic genes. Biological Psychiatry, 83(5):456–465.

McKinney, W. et al. (2010). Data structures for statistical computing in Python. In Proceedings of the 9th Python in Science Conference, volume 445, pages 51–56. Austin, TX.

Messé, A. (2020). Parcellation influence on the connectivity-based structure-function relationship in the human brain. Human Brain Mapping, 41(5):1167–1180.

Miller, J. A., Ding, S.-L., Sunkin, S. M., Smith, K. A., Ng, L., Szafer, A., Ebbert, A., Riley, Z. L., Royall, J. J., Aiona, K., et al. (2014). Transcriptional landscape of the prenatal human brain. Nature, 508(7495):199–206.

Mišić, B., Fatima, Z., Askren, M. K., Buschkuehl, M., Churchill, N., Cimprich, B., Deldin, P. J., Jaeggi, S., Jung, M., Korostil, M., et al. (2014). The functional connectivity landscape of the human brain. PLoS One, 9(10):e111007.

Morgan, S. E., Seidlitz, J., Whitaker, K. J., Romero-Garcia, R., Clifton, N. E., Scarpazza, C., van Amelsvoort, T., Marcelis, M., van Os, J., Donohoe, G., et al. (2019). Cortical patterning of abnormal morphometric similarity in psychosis is associated with brain expression of schizophrenia-related genes. Proceedings of the National Academy of Sciences, 116(19):9604–9609.

Negi, S. K. and Guda, C. (2017). Global gene expression profiling of healthy human brain and its application in studying neurological disorders. Scientific Reports, 7(1):1–12.

Nørgaard, M., Beliveau, V., Ganz, M., Svarer, C., Pinborg, L. H., Keller, S. H., Jensen, P. S., Greve, D. N., and Knudsen, G. M. (2021). A high-resolution in vivo atlas of the human brain’s benzodiazepine binding site of GABA_A_ receptors. NeuroImage, 232:117878.

Oldham, M. C., Konopka, G., Iwamoto, K., Langfelder, P., Kato, T., Horvath, S., and Geschwind, D. H. (2008). Functional organization of the transcriptome in human brain. Nature Neuroscience, 11(11):1271.

Oldham, S., Arnatkevičiūtė, A., Smith, R. E., Tiego, J., Bellgrove, M. A., and Fornito, A. (2020). The efficacy of different preprocessing steps in reducing motion-related confounds in diffusion MRI connectomics. NeuroImage, 222:117252.

Oliphant, T. E. (2006). A guide to NumPy, volume 1. Trelgol Publishing USA.

Park, B.-y., Bethlehem, R. A., Paquola, C., Larivière, S., Rodríguez-Cruces, R., de Wael, R. V., the NSPN consortium, Bullmore, E. T., and Bernhardt, B. C. (2021). An expanding manifold in transmodal regions characterizes adolescent reconfiguration of structural connectome organization. Elife, 10:e64694.

Park, B.-y., Park, H., Morys, F., Kim, M., Byeon, K., Lee, H., Kim, S.-H., Valk, S., Dagher, A., and Bernhardt, B. (2020). Body mass variations relate to fractionated functional brain hierarchies. BioRxiv.

Parkes, L., Fulcher, B., Yücel, M., and Fornito, A. (2017). Transcriptional signatures of connectomic subregions of the human striatum. Genes, Brain and Behavior, 16(7):647–663.

Parkes, L., Fulcher, B., Yücel, M., and Fornito, A. (2018). An evaluation of the efficacy, reliability, and sensitivity of motion correction strategies for resting-state functional MRI. NeuroImage, 171:415–436.

Pedregosa, F., Varoquaux, G., Gramfort, A., Michel, V., Thirion, B., Grisel, O., Blondel, M., Prettenhofer, P., Weiss, R., Dubourg, V., et al. (2011). Scikit-learn: Machine learning in Python. Journal of Machine Learning Research, 12(Oct):2825–2830.

Pérez, F. and Granger, B. E. (2007). IPython: A system for interactive scientific computing. Computing in Science & Engineering, 9(3):21–29.

Preller, K. H., Burt, J. B., Ji, J. L., Schleifer, C. H., Adkinson, B. D., Stämpfli, P., Seifritz, E., Repovs, G., Krystal, J. H., Murray, J. D., et al. (2018). Changes in global and thalamic brain connectivity in LSD-induced altered states of consciousness are attributable to the 5-HT2A receptor. Elife, 7:e35082.

Richiardi, J., Altmann, A., and Greicius, M. (2017). Distance is not everything in imaging genomics of functional networks: reply to a commentary on *Correlated gene expression supports synchronous activity in brain networks*. bioRxiv, page 132746.

Richiardi, J., Altmann, A., Milazzo, A.-C., Chang, C., Chakravarty, M. M., Banaschewski, T., Barker, G. J., Bokde, A. L., Bromberg, U., Büchel, C., et al. (2015). Correlated gene expression supports synchronous activity in brain networks. Science, 348(6240):1241–1244.

Rittman, T., Rittman, M., and Azevedo, T. (2017). Maybrain software package. https://github.com/RittmanResearch/maybrain.

Rittman, T., Rubinov, M., Vértes, P. E., Patel, A. X., Ginestet, C. E., Ghosh, B. C., Barker, R. A., Spillantini, M. G., Bullmore, E. T., and Rowe, J. B. (2016). Regional expression of the MAPT gene is associated with loss of hubs in brain networks and cognitive impairment in parkinson disease and progressive supranuclear palsy. Neurobiology of Aging, 48:153–160.

Rizzo, G., Veronese, M., Expert, P., Turkheimer, F. E., and Bertoldo, A. (2016). MENGA: a new comprehensive tool for the integration of neuroimaging data and the Allen Human Brain transcriptome atlas. PloS one, 11(2):e0148744.

Roberts, J. A., Perry, A., Lord, A. R., Roberts, G., Mitchell, P. B., Smith, R. E., Calamante, F., and Breakspear, M. (2016). The contribution of geometry to the human connectome. Neuroimage, 124:379–393.

Romero-Garcia, R., Whitaker, K. J., Váša, F., Seidlitz, J., Shinn, M., Fonagy, P., Dolan, R. J., Jones, P. B., Goodyer, I. M., Bullmore, E. T., et al. (2018). Structural covariance networks are coupled to expression of genes enriched in supragranular layers of the human cortex. NeuroImage, 171:256–267.

Romme, I. A., de Reus, M. A., Ophoff, R. A., Kahn, R. S., and van den Heuvel, M. P. (2017). Connectome disconnectivity and cortical gene expression in patients with schizophrenia. Biological Psychiatry, 81(6):495–502.

Rousseeuw, P. J. (1987). Silhouettes: a graphical aid to the interpretation and validation of cluster analysis. Journal of Computational and Applied Mathematics, 20:53–65.

Schilling, K. G., Nath, V., Hansen, C., Parvathaneni, P., Blaber, J., Gao, Y., Neher, P., Aydogan, D. B., Shi, Y., Ocampo-Pineda, M., et al. (2019). Limits to anatomical accuracy of diffusion tractography using modern approaches. NeuroImage, 185:1–11.

Seidlitz, J., Nadig, A., Liu, S., Bethlehem, R. A. I., Vértes, P. E., Morgan, S. E., Váša, F., Romero-Garcia, R., Lalonde, F. M., Clasen, L. S., Blumenthal, J. D., Paquola, C., Bernhardt, B., Wagstyl, K., Polioudakis, D., de la Torre-Ubieta, L., Geschwind, D. H., Han, J. C., Lee, N. R., Murphy, D. G., Bullmore, E. T., and Raznahan, A. (2020). Transcriptomic and cellular decoding of regional brain vulnerability to neurogenetic disorders. Nature Communications, 11(1):1–14.

Seidlitz, J., Váša, F., Shinn, M., Romero-Garcia, R., Whitaker, K. J., Vértes, P. E., Wagstyl, K., Reardon, P. K., Clasen, L., Liu, S., Messinger, A., Leopold, D. A., Fonagy, P., Dolan, R. J., Jones, P. B., Goodyer, I. M., the NSPN Consortium, Raznahan, A., and T.Bullmore, E. (2018). Morphometric similarity networks detect microscale cortical organization and predict interindividual cognitive variation. Neuron, 97(1):231–247.

Sepulcre, J., Grothe, M. J., Uquillas, F. d., Ortiz-Terán, L., Diez, I., Yang, H.-S., Jacobs, H. I., Hanseeuw, B. J., Li, Q., El-Fakhri, G., et al. (2018). Neurogenetic contributions to amyloid beta and tau spreading in the human cortex. Nature Medicine, 24(12):1910–1918.

Shafiei, G., Bazinet, V., Dadar, M., Manera, A. L., Collins, D. L., Dagher, A., authors, G., Misic, B., and Ducharme, S. (2021). Atrophy patterns in sporadic and genetic behavioral variant frontotemporal dementia are based on brain network architecture. bioRxiv.

Shafiei, G., Markello, R. D., De Wael, R. V., Bernhardt, B. C., Fulcher, B. D., and Misic, B. (2020). Topographic gradients of intrinsic dynamics across neocortex. Elife, 9:e62116.

Shin, J., French, L., Xu, T., Leonard, G., Perron, M., Pike, G. B., Richer, L., Veillette, S., Pausova, Z., and Paus, T. (2018). Cellspecific gene-expression profiles and cortical thickness in the human brain. Cerebral Cortex, 28(9):3267–3277.

Shine, J. M., Breakspear, M., Bell, P. T., Martens, K. A. E., Shine, R., Koyejo, O., Sporns, O., and Poldrack, R. A. (2019). Human cognition involves the dynamic integration of neural activity and neuromodulatory systems. Nature Neuroscience, 22(2):289–296.

Simmons, J. P., Nelson, L. D., and Simonsohn, U. (2011). Falsepositive psychology: Undisclosed flexibility in data collection and analysis allows presenting anything as significant. Psychological Science, 22(11):1359–1366.

Sousa, A. M., Zhu, Y., Raghanti, M. A., Kitchen, R. R., Onorati, M., Tebbenkamp, A. T., Stutz, B., Meyer, K. A., Li, M., Kawasawa, Y. I., et al. (2017). Molecular and cellular reorganization of neural circuits in the human lineage. Science, 358(6366):1027–1032.

Steegen, S., Tuerlinckx, F., Gelman, A., and Vanpaemel, W. (2016). Increasing transparency through a multiverse analysis. Perspectives on Psychological Science, 11(5):702–712.

Thirion, B., Varoquaux, G., Dohmatob, E., and Poline, J.-B. (2014). Which fMRI clustering gives good brain parcellations? Frontiers in Neuroscience, 8:167.

Thompson, W. H., Wright, J., Bissett, P. G., and Poldrack, R. A. (2020). Meta-Research: Dataset decay and the problem of sequential analyses on open datasets. ELife, 9:e53498.

Valk, S. L., Kanske, P., Park, B.-y., Hong, S. J., Boeckler-Raettig, A., Trautwein, F.-M., Bernhardt, B. C., and Singer, T. (2021). Functional network plasticity of the human social brain. bioRxiv, pages 2020–11.

Van Der Walt, S., Colbert, S. C., and Varoquaux, G. (2011). The NumPy array: a structure for efficient numerical computation. Computing in Science & Engineering, 13(2):22.

Van Essen, D. C., Smith, S. M., Barch, D. M., Behrens, T. E., Yacoub, E., Ugurbil, K., Consortium, W.-M. H., et al. (2013). The WU-Minn human connectome project: an overview. NeuroImage, 80:62–79.

Vértes, P. E., Rittman, T., Whitaker, K. J., Romero-Garcia, R., Váša, F., Kitzbichler, M. G., Wagstyl, K., Fonagy, P., Dolan, R. J., Jones, P. B., et al. (2016). Gene transcription profiles associated with inter-modular hubs and connection distance in human functional magnetic resonance imaging networks. Philosophical Transactions of the Royal Society B: Biological Sciences, 371(1705):20150362.

Virtanen, P., Gommers, R., Oliphant, T. E., Haberland, M., Reddy, T., Cournapeau, D., Burovski, E., Peterson, P., Weckesser, W., Bright, J., et al. (2020). SciPy 1.0: fundamental algorithms for scientific computing in Python. Nature Methods, pages 1–12.

Vogel, J. W., Iturria-Medina, Y., Strandberg, O. T., Smith, R., Levitis, E., Evans, A. C., and Hansson, O. (2020). Spread of pathological tau proteins through communicating neurons in human alzheimer’s disease. Nature Communications, 11(1):1–15.

Wang, D., Liu, S., Warrell, J., Won, H., Shi, X., Navarro, F. C., Clarke, D., Gu, M., Emani, P., Yang, Y. T., et al. (2018). Comprehensive functional genomic resource and integrative model for the human brain. Science, 362(6420).

Waskom, M., Botvinnik, O., O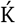ane, D., Hobson, P., Ostblom, J., Lukauskas, S., Gemperline, D. C., Augspurger, T., Halchenko, Y., Cole, J. B., and et al. (2018). mwaskom/seaborn. Zenodo, doi:10.5281/zenodo.592845.

Waskom, M., Larson, E., Brodbeck, C., Gramfort, A., Burns, S., Luessi, M., Weidemann, C. T., Bitzer, S., Markiewicz, C., LaPlante, R., Halchenko, Y., Engemann, D. A., van Vliet, M., Ghosh, S., Klein, N., Piantoni, G., Brett, M., Gwilliams, L., King, J.-R., and Liu, D. (2020). nipy/pysurfer. Zenodo, doi:10.5281/zenodo.592515.

Whitaker, K. J., Vértes, P. E., Romero-Garcia, R., Váša, F., Moutoussis, M., Prabhu, G., Weiskopf, N., Callaghan, M. F., Wagstyl, K., Rittman, T., Tait, R., Ooi, C., Suckling, J., Inkster, B., Fonagy, P., Dolan, R. J., Jones, P. B., Goodyer, I. M., the NSPN Consortium, and Bullmore, E. T. (2016). Adolescence is associated with genomically patterned consolidation of the hubs of the human brain connectome. Proceedings of the National Academy of Sciences, 113(32):9105–9110.

Yao, Z., van Velthoven, C. T., Nguyen, T. N., Goldy, J., Sedeno-Cortes, A. E., Baftizadeh, F., Bertagnolli, D., Casper, T., Chiang, M., Crichton, K., et al. (2021). A taxonomy of transcriptomic cell types across the isocortex and hippocampal formation. Cell, 184(12):3222–3241.

Zhao, K., Zheng, Q., Che, T., Martin, D., Li, Q., Ding, Y., Zheng, Y., Liu, Y., and Li, S. (2020). Regional radiomics similarity networks (R2SN) in the human brain: reproducibility, smallworld and biological basis. bioRxiv.

Zheng, Y.-Q., Zhang, Y., Yau, Y., Zeighami, Y., Larcher, K., Misic, B., and Dagher, A. (2019). Local vulnerability and global connectivity jointly shape neurodegenerative disease propagation. PLoS biology, 17(11):e3000495.

